# Distinct Roles of CaMKII in Synaptic Vesicle Dynamics at Zebrafish Retinal Rod Bipolar Ribbon Synapses

**DOI:** 10.1101/2025.07.27.667036

**Authors:** Johane M. Boff, Nirujan Rameshkumar, Moumita Khamrai, Jayaraman Seetharaman, Cindy Lorena Olmos-Carreño, Nabeel Chudasama, Takeshi Yoshimatsu, David Zenisek, Steven J. Tavalin, Thirumalini Vaithianathan

**Affiliations:** Department of Pharmacology, Addiction Science, and Toxicology, College of Medicine, University of Tennessee Health Science Center, Memphis, TN 38163, USA; Department of Ophthalmology and Visual Sciences, Washington University in St Louis School of Medicine, St. Louis, MO 63110, USA; Department of Health, University of Waterloo, Waterloo, ON N2L3G5, Canada; Department of Cellular and Molecular Physiology, Yale University School of Medicine, New Haven, CT 06520-8066, USA; Department of Ophthalmology and Visual Sciences, Yale University School of Medicine, New Haven, CT 06520-8066, USA; Department of Ophthalmology, University of Tennessee Health Science Center, Memphis, TN 38163, USA

## Abstract

Calcium (Ca²⁺) not only serves as a fundamental trigger for neurotransmitter release but also participates in shaping neurotransmitter release (NTR) during prolonged presynaptic stimulation via multiple Ca^2+^-dependent processes. The Ca^2+^/calmodulin (CaM)-dependent protein kinase II (CaMKII) is enriched at various presynaptic terminals, including ribbon synapses, where it associates with synaptic ribbons and thus may contribute to the modulation of Ca^2+^-dependent NTR. This could arise via its ability to influence one or more steps that control either the Ca^2+^ signal, the release process, or synaptic vesicle dynamics affecting available pools. Yet, recent studies have yielded conflicting results regarding the ability of CaMKII to influence NTR at rod bipolar cell (RBC) ribbon synapses. To address this, we acutely manipulated CaMKII activity in synaptic terminals of zebrafish RBCs by infusion of either inhibitory peptides targeting CaMKII or CaM, or a constitutively active CaMKII, while using a combination of imaging and electrophysiological approaches. Neither inhibiting nor enhancing CaMKII activity affects presynaptic Ca^2+^ channel activity. However, capacitance measurements revealed that inhibition of either CaMKII or CaM reduces exocytosis. CaMKII inhibition also reduces synaptic vesicle replenishment. Surprisingly, elevation of CaMKII activity also diminished vesicle fusion, similar to the effect of CaMKII inhibition, suggesting that CaMKII activity naturally exists at optimal levels to support neurotransmitter release. In contrast to CaMKII inhibition, CaMKII activity elevation did not impair vesicle replenishment. Collectively, these data suggest that distinct synaptic vesicle populations are differentially reliant on the level of CaMKII activity.

**Significant Statement:** CaMKII is well-recognized for its postsynaptic participation in multiple forms of synaptic plasticity, yet it is well-documented to be prevalent in presynaptic compartments, where it may control NTR. The specific functions of CaMKII in synaptic vesicle dynamics remain poorly understood. While using a combination of imaging and electrophysiology approaches, we acutely manipulated CaMKII activity in presynaptic terminals by infusion of inhibitory peptides or constitutively active CaMKII. These manipulations revealed that deviations in CaMKII levels, in either direction, impair neurotransmitter release, suggesting that CaMKII is optimally present at these synapses. Yet CaMKII activity appears required for synaptic vesicle replenishment, suggesting that distinct aspects of synaptic vesicle dynamics are under differential control by CaMKII.

## Introduction

NTR is tightly regulated by a complex interplay of cellular mechanisms, including the dynamic balance between exocytosis and vesicle replenishment at release sites. Sensory neurons such as photoreceptors, bipolar cells, and hair cells contain specialized ribbon synapses, where local Ca^2+^ signals control kinetically distinct phases of NTR (Mennerick and Matthews, 1996; Beutner and Moser, 2001; Rodriguez-Contreras and Yamoah, 2001; Singer and Diamond, 2006; Babai, Bartoletti and Thoreson, 2010; Babai, Morgans and Thoreson, 2010; Vaithianathan and Matthews, 2014). CaM regulates Ca²⁺-dependent vesicle recruitment and exocytosis at both conventional synapses (Peters and Mayer, 1998; Persechini and Cronk, 1999; Quetglas et al., 2000; Sakaba and Neher, 2001; Lipstein et al., 2013; Timofeeva and Volynski, 2015) and cone photoreceptors (Van Hook et al., 2014). CaMKII, a downstream effector of CaM, is highly enriched at presynaptic terminals (Gorelick et al., 1988; Walaas, Gorelick and Greengard, 1989) and associates with synaptic vesicles and ribbon synapses (Takamori et al., 2006; Uthaiah and Hudspeth, 2010b; Kantardzhieva et al., 2012). Furthermore, in rod bipolar cells (RBCs), CaMKII phosphorylates syntaxin-3B, a SNARE protein linked to retinal neurotransmission, in a Ca^2+^-dependent manner (Liu, Heidelberger and Janz, 2014; Campbell et al., 2020). However, CaMKII’s role on NTR in both central and ribbon synapses is unclear, as previous work demonstrated that CaMKII can increase (Jin and Hawkins, 2003; Lu and Hawkins, 2006; Liu, Heidelberger and Janz, 2014; Campbell et al., 2020), decrease (Hinds et al., 2003), or even have minimal to no effect (Liang et al., 2021), in part due to different experimental conditions and synapses being investigated (Wang, 2008). Given that CaMKII is present within the ribbon complex (Uthaiah and Hudspeth, 2010a), it seems plausible that CaMKII activity may influence various aspects of NTR at ribbon synapses.

CaMKII consists of four isoforms (α, β, γ, and δ) encoded by different genes that primarily differ in the length and composition of their linker domains, which contribute to isoform-specific localization and function (Bhattacharyya et al., 2020). In the mammalian retina, these isoforms display overlapping but distinct expression patterns (Terashima, Ochiishi and Yamauchi, 1994; Tetenborg et al., 2017), with CaMKIIγ exhibiting diffuse expression across the retina and CaMKIIδ being particularly prominent in RBC soma and terminals, suggesting these isoforms are likely the most relevant for these cells. Although the functions of these isoforms have not been thoroughly investigated in the retina, their distinct expression patterns suggest that each isoform may play specialized roles in modulating retinal circuitry and visual processing.

CaMKII participation in retinal transmission has been commonly probed using CaMKII inhibitors including KN-62, KN-93, and various inhibitory peptides derived from the autoinhibitory domain of CaMKII. Yet, substantial off-target effects of KN-62 and KN-93 have been noted (Brown and Bayer, 2024) and thus use of autoinhibitory domain-derived peptides such as autocamtide-2-related inhibitory peptide (AIP) or autocamtide-3-derived peptide (AC3-I) may be preferred due to their enhanced potency and/or specificity. Yet, peptides derived from this CaMKII region may also exhibit off-target effects (Hvalby et al., 1994; Backs et al., 2009). As such, in this study, we infused an optimized peptide (CN19o) derived from the endogenous CaMKII inhibitor protein (CaMKIIN), which may offer improved potency and selectivity (Brown and Bayer, 2024), to acutely examine the contribution of endogenous CaMKII to NTR. Complementary experiments in which CaMKII levels were acutely elevated by infusion of exogenous constitutively active (CA) CaMKII provided an additional means to evaluate the relative range in which CaMKII operates to control various aspects of NTR release. These studies reveal that CaMKII likely already exists at optimal levels in zebrafish RBCs to support release and that deviations in CaMKII activity, in either direction, impair NTR. However, CaMKII activity appears to be required for the recovery of the releasable pool of synaptic vesicles following a sustained depolarization.

## Materials and Methods

### Zebrafish Rearing

Adult zebrafish were used for all experiments, with specific age ranges and strains depending on the technique. All animals were euthanized in ice-chilled conditioned water, and retinas or eyecups were dissected as required. For electrophysiological recordings, male and female zebrafish aged between 14 and 20 months were raised under a 14-hour light/10-hour dark cycle. The zebrafish were dark-adapted to allow the separation of the retinal pigment epithelium from the retina. In-Situ Hybridization Chain Reaction was performed on adult (9-12 months old) *Tg(vsx1:mem-Cerulean)* transgenic fish, which express membrane-targeted Cerulean fluorescent protein in RBC1 (Hellevik et al., 2024). For Serial Block-Face Scanning Electron Microscopy (SBF-SEM), eyecups were dissected in bicarbonate Ames’ medium and fixed in 4% glutaraldehyde in 0.1M sodium-cacodylate buffer. All animal procedures were performed in agreement with the University of Tennessee Health Science Center Guidelines for Animals in Research and Washington University in St. Louis, Institutional Animal Care and Use Committee (IACUC) guidelines.

### RBC Isolation

Retinal bipolar cells were isolated as described previously (Vaithianathan and Matthews, 2014; Vaithianathan et al., 2019; Shrestha and Vaithianathan, 2022; Shrestha et al., 2023). Dark-adapted zebrafish were anesthetized in cold 4°C conditioned water for 20 minutes and euthanized by quick decapitation. After the removal of both eyes and retinal extraction, retinal tissue was treated in an oxygenated saline solution containing 115 mM NaCl, 2.5 mM KCl, 0.5 mM CaCl_2_, 1 mM MgCl_2_, 10 mM HEPES, 10 mM glucose, pH 7.4, supplemented with 1,100 units/mL type V hyaluronidase (Worthington Biochemical Corp., Lakewood, NJ, USA) at 25°C for 25 minutes. Retinal pieces were washed with saline and incubated in a mixture of saline with 5 mM DL-cysteine and 20-30 units/mL papain (Sigma-Aldrich, St. Louis, MO, USA). The tissue was triturated using a fire-polished glass Pasteur pipette, and the dissociated cells were plated onto glass-bottomed dishes, prepared as previously described (Shrestha and Vaithianathan, 2022), in saline. After allowing the cells to settle on the dish for 30 minutes, the saline solution was exchanged for another containing 2.5 mM CaCl_2_. RBCs used for experiments were identified based on their characteristic morphology (Hellevik et al., 2024).

### CaMKII reagent synthesis and validation

#### CN19o

An optimized form of CaMKIINtide (CN19o: KRAPKLGQIGRQKAVDIED) (Coultrap and Bayer, 2011) and a scrambled form (CN19scr: VKEARIDGKPDRLAGQKQI) were synthesized (LifeTein) with a single cysteine residue added on the N-terminal side that was conjugated with Cy5. These peptides were evaluated for their ability to inhibit the α, γ, and δ isoforms of GST-tagged full-length human CaMKII (Promega; 10 ng). Ca^2+^ (400 µM)/CaM (5 ng/µl)-dependent activity was evaluated using autocamtide-2 (130 µM) as substrate in reaction buffer (final concentrations: 20 mM Tris-HCl, 10 mM MgCl_2_, 50 μg/ml BSA; pH 7.4). Peptides were added to the kinase/substrate reaction 5 min prior to initiation. Reactions (50 µl) were initiated with the addition of ATP (to a final concentration of 100 µM) and carried out at 30°C for 5, 10, or 30 minutes for CaMKII8, CaMKIIγ, or CaMKIIα, respectively, upon which they were terminated by the addition of 50 µl ATP-glo substrate (Promega). Luminescence was acquired using a Bio-Rad XRS chemiluminescence documentation system and Quantity One software. Signals from each reaction were compared against a standard curve (0-100 µM ATP) to derive the amount of ATP consumed during the reaction. ATP consumed in the presence of CN19o and CN19o scrambled (1 μm) was normalized to that obtained in the absence of peptides. The resulting curves were fit with a logistic equation using OriginPro software (version 9.1).

### CaM binding domain (CaMBD)

An N-terminal His-tagged fragment of the rat SK2 channel (residues 412-487) in pET33b provided by John Adelman (Vollum Institute), designated here as CaMBD, was transformed into BL21(DE3) cells (Novagen) and grown overnight in LB media (200 mL) supplemented with kanamycin. Cells were harvested by centrifugation and purified using the Ni-NTA fast start kit (Qiagen) as previously described (Brooks and Tavalin, 2011). The resulting protein was dialyzed three times in PBS supplemented with 10% glycerol (pH 7.40; Invitrogen) using a 3.5 kDa MWCO dialysis cassette (ThermoFisher Scientific) and then concentrated using Amicon Ultra 3 kDa MWCO centrifugal filter units (MilliporeSigma). Protein concentration was determined with a modified Bradford assay (Bio-Rad) using bovine serum albumin (BSA) as standard. The concentration of protein was adjusted to 5 ug/µl (500 µM) with PBS and aliquoted in convenient volumes for eventual addition to the patch pipette solution. To evaluate Ca^2+^-independent binding of CaM to CaMBD, CaMBD (5 μg) was incubated with 10 µl His-tag magnetic beads (Invitrogen) for 30 minutes in 500 µL of IP buffer (150 mM NaCl, 10 mM HEPES, 1 mM EGTA, 0.1% Tween-20; pH 7.40) and then washed three times to remove unbound CaMBD. CaM (∼85 µg) was added to this buffer to achieve a concentration of 10 µM and incubated overnight at 4°C. Beads were washed three times (5 min each) and then resuspended in 20 µL of SDS sample buffer, boiled for 5 minutes, and resolved by SDS-PAGE. Blots were probed with a monoclonal antibody to CaM (1:1000 dilution; Millipore) followed by an HRP-conjugated goat anti-mouse secondary antibody (MilliporeSigma) and visualized by enhanced chemiluminescence using a Bio-Rad Chemidoc XRS imaging system. Blots were stripped and then probed with an HRP-conjugated mouse monoclonal anti-His antibody (1:5,000 dilution) and visualized to verify the presence of the His-tagged CaMBD.

### CA His-CaMKII(1-290)

CA His-CaMKII(1-290) in pET100/D-TOPO was previously described (Tavalin, 2008) and produced as recently described (Tavalin and Colbran, 2017) by transformation into Rosetta2(DE3)pLysS cells (Novagen) and grown overnight at 30°C in LB media (10 mL) supplemented with 100 µg/mL carbenicillin. This starter culture was diluted to 250 mL and grown to an OD_600_ ∼0.6 and then induced with IPTG (1 mM) for 3 hours. Cells were harvested by centrifugation and then lysed in B-PER (ThermoFisher Scientific) and purified using the HisPur Cobalt purificiation kit (ThermoFisher Scientific). The resulting protein was dialyzed three times in kinase storage buffer (100 mM NaCl, 20 mM HEPES, 0.1 mM EDTA, 2 mM DTT, and 10% glycerol; pH 7.4) using a 10 kDa MWCO dialysis cassette (ThermoFisher Scientific) and then concentrated using Amicon Ultra 10 kDa MWCO centrifugal filter units (MilliporeSigma). Protein concentration was determined with a modified Bradford assay (Bio-Rad) using bovine serum albumin (BSA) as standard. The concentration of protein was adjusted to 740 ng/µL (20 µM) with kinase storage buffer and aliquoted in convenient volumes for eventual addition to the patch pipette solution. The kinase was heat inactivated by incubating at 95 °C for 20 minutes. Ca^2+^/CaM-independent constitutive activity of the kinase was assayed in 50 µL reactions and initiated and terminated similarly to that described above, using phosphorylation buffer (10 mM HEPES, 10 mM MgCl2, 1 mM DTT, and 0.1 mg/ml BSA, pH 7.40) and syntide (100 µM) as substrate. The specific activity of the kinase (at 100 ng) was 810 ± 81 nmol/min/mg.

### RBC Voltage Patch-Clamp Recordings

Whole-cell recordings were obtained by placing the patch pipette (7-12 MΩ) at the terminals of RBCs. The patch pipette solution was composed of 120 mM Cs-gluconate, 20 mM HEPES, 10 mM tetraethylammonium chloride, 3 mM MgCl_2_, 0.2 mM N-methyl-D-glucamine (NMDG)-EGTA, 2 mM Mg-ATP, and 0.5 mM GTP. The pipette solution was either used by itself or mixed with CN19o (1 μM), scrambled CN19o (CN19scr; 1 μM), a calmodulin-binding domain (CaMBD) (10 μM), constitutively active (CA) His-CaMKII(1-290) (0.1, 10, or 100 nM), or its heat-inactivated (HI) version (100 nM) depending on the experimental conditions. The extracellular solution was the saline solution described above, containing 2.5 mM CaCl_2_. All measurements were taken 5 minutes after establishing whole-cell configuration, as the average dye loading rate was approximately 1.27 minutes. This rate was determined by performing xyt scans to monitor fluorescence over time within a defined region of interest (3×3 pixels). Fluorescence intensity data were fitted using a sigmoid function to extract the loading kinetics, yielding an average loading rate of 1.27 minutes for the peptide. Ca^2+^ currents were evoked under the voltage-clamp using a HEKA EPC-10 amplifier controlled by PatchMaster software (HEKA Instruments, Inc., Holliston, MA, USA; version v2×90.4) with a holding potential of-65 mV and stepped to 0 mV for different lengths depending on experimental goals. Currents were acquired at 50 kHz and filtered at 2.9 kHz. For IV experiments, 9 voltage steps were applied with a 250 ms duration, and P/4 leak subtraction was applied by the Patchmaster software by applying four leak pulses with a 100 ms delay prior to test depolarization. Readings from the same cell at-10 mV were averaged for statistical analysis. Membrane capacitance was measured using the sine DC method of the PatchMaster Lock-in extension. Only the first sustained pulse recording was used for capacitance data analysis with a series resistance smaller than ∼ 75 MΩ.

### Laser-Scanning Confocal Microscopy

All fluorescence imaging was performed using a 60x silicon objective on an Olympus FV-3000 laser-scanning confocal microscopy system (Olympus, Shinjuku, Tokyo, Japan) controlled by the Olympus FV31S0SW software (version 2.3.1.163) with a Galvano scanner. Acquisition parameters such as pinhole diameter, laser power, PMT gain, scan speed, optical zoom, offset, and step size were kept constant between experiments. To localize synaptic ribbons and measure local Ca^2+^ signals, the pipette solution was supplemented with 35 µM 5-carboxy tetramethylrhodamine (TAMRA: GIDEEKPVDLTAGRRAG)-labeled ribbon-binding peptides (RBP) (LifeTein, LLC, Somerset, NJ, USA) and a custom-made cysteine-containing ribeye binding peptide NH_2_-CIEDEEKPVDLTAGRRAC-COOH (LifeTein, LLC, Somerset, NJ, USA), which was conjugated with fluorogenic 520^®^ maleimide low-affinity, here referred to as Cal520L (*K_D_*, 90 µM, purchased from AAT Bioquest, CA) (Rameshkumar et al., 2024). Depending on the experiment, this was also combined with 1 μM CN19o. Sequential line scans were acquired at 1.51 ms/line and 10.0 μs/pixel with a scan size of 256×256 pixels. Image acquisition timing was established via transistor-transistor (TTL) pulses between the FluoView software (version 2.3.1.163) and the PatchMaster software (version v2×90.4) used for patch-clamping (HEKA), ensuring synchronization of voltage-clamp and imaging data

### Single-Cell RNA-Sequencing

We retrieved pre-processed count matrices from published studies for adult zebrafish bipolar cells (Hellevik et al., 2024) and replotted the average and percent expressions for the genes of interest.

### In-situ Hybridization Chain Reaction

Adult (9-12 months old) *Tg(vsx1:mem-Cerulean)* transgenic fish, which express membrane-targeted Cerulean fluorescent protein in RBC1 (Hellevik et al., 2024). Adult zebrafish were humanely euthanized in ice-chilled fish water. After decapitation, retinal tissues were dissected from the enucleated whole eyes by removing cornea, lens, and epithelial layer in 1x PBS. The tissues were immediately fixed in 4% paraformaldehyde (Agar Scientific, AGR1026) in PBS for 30 min at room temperature, followed by three washes in PBS. The tissues were then sliced at 200 μm thickness using a tissue chopper. The standard in situ HCR was performed according to the manufacturer’s protocol using HCR Probe hybridization buffer, Probe Wash buffer, and Amplification buffer (Molecular Instruments). HCR probe sets and Amplifiers were custom-designed (Table 1). Confocal image stacks were taken immediately after the in-situ HCR on a FV1000 microscope (Olympus) with a 40×780 oil immersion objective (HC PL APO CS2, Leica). Typical voxel size was 0.62 μm and 0.5 μm in the 𝑥-𝑦 and 𝑧, respectively. Contrast, brightness, and pseudo-color were adjusted for display in FIJI. Image analysis and quantification were performed using FIJI. For each RBC1, a region of interest was drawn around the soma, and a subset of z-slices was selected where the cell was present. A binary mask of the RBC was generated by thresholding the Cerulean fluorescence channel, and the in situ channel was isolated in a separate task. The mask was used to obtain the number of pixels inside the cell and then multiplied by the signal from the situ channel. This was then z-projected, and both the area and mean intensity of the signal were measured. Background measurements were obtained from a blank region and multiplied by the number of pixels inside the cell to match the volume of the cell stack. The final normalized in situ signal for each cell was calculated using the formula:

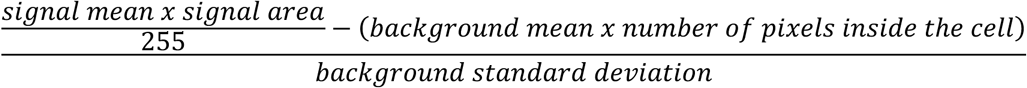

### Serial Block-Face Scanning Electron Microscopy (SBF-SEM)

Zebrafish eyecups were dissected in bicarbonate Ames’ medium and fixed in 4% glutaraldehyde in 0.1M sodium-cacodylate buffer at 25°C for an hour and at 4°C overnight. The eyecups were then treated for SBF-SEM in Zeiss 3-View SBF-SEM, as described previously (Della Santina et al., 2016). Montages were acquired throughout the entire retina and each montage was 35 μm long, with a 5 nm XY resolution and 50 nm Z resolution. The montages were stitched together, processed, viewed, and analyzed using ImageJ’s TrakEM2 plugin (version 1.5h, NIH). RBCs were identified based on their characteristic morphology and size (Hellevik et al., 2024; Rameshkumar et al., 2024) and the number of vesicles was counted for each cell’s ribbons and averaged.

### Data Processing

Data was obtained from FluoView and PatchMaster and exported to Igor Pro (Version 8.04), Excel (Version 16.76), and/or ImageJ (imagej.nih.gov) for further analysis and image production, and RStudio (version 2023.09.0+463) software for statistical analysis and scRNA-seq data processing and image production. Figures were assembled using Adobe Photoshop (Version 25.12.0).

### Experimental Design and Statistical Analysis

The design of each experiment is described above. Between-group differences were analyzed using unpaired t-tests, with Welch’s correction applied when variances were unequal as determined by an F-test. All statistical analyses were performed using RStudio (version 2023.09.0+463). Data are represented as mean ± SEM unless otherwise noted in figure legends. Sample size for each experiment is reported within the results section and in the figure legends.

### Data and Code Accessibility

The datasets used in this study will be provided by the corresponding author upon reasonable request. All RStudio codes used for data processing, statistical analysis, and scRNA-seq figure generation are available on GitHub (https://github.com/vaithianathanlab/camkii_manuscript_codes).

## Results

### CaMKII isoform prevalence in rod bipolar cells

In the mouse retina, CaMKIIγ is diffusely expressed in the entire retina, and CaMKIIδ is more specifically present in RBC terminals (Tetenborg et al., 2017), whereas in the rat retina, α and β isoforms of CaMKII are prevalently expressed in most ganglion cells (Terashima, Ochiishi and Yamauchi, 1994), suggesting CaMKII expression in the retina is likely to be species-dependent. To gain insight into the distribution of CaMKII isoforms in the zebrafish retina, we first analyzed the published scRNA-seq dataset (GEO accession number: GSE237214) (Hellevik et al., 2024). This study reported that the zebrafish retina contains two types of RBCs, RBC1 and RBC2, which can be identified based on their different morphologies, *prkca* expression (high in RBC1) (Supplementary Fig. 1), and specific labeling in different transgenic lines: RBC1 in *Tg(vsx1:mem-Cerulean)* and RBC2 in *Tg(vsx2:mem-Cerulean)*. RBC1 best resembles the mammalian RBCs, participating in synapses with narrow-field A2 and wide-field A17 amacrine cells, whereas RBC2 exclusively synapses with wide-field amacrine cells (Hellevik et al., 2024). Here, we focus on zebrafish RBC1s, which we will refer to as “RBCs” for simplicity. Based on this data, CaMKIIγ is the predominant isoform in both types of RBCs (Supplementary Fig. 1A), followed by the δ isoform, then the β, and with minimal levels of the α. In addition to CaMKII, we also analyzed the prevalence of the isoforms of many other direct and indirect downstream targets of CaM, including myosin light-chain kinase (MYLK), phosphodiesterase 1 (PDE1), and AMP-activated protein kinase (AMPK) (Supplementary Fig. 1A). Of these, CaMKII is the highest expressed Ca^2+^/CaM-dependent signaling enzymes in RBCs, consistent with its colocalization with synaptic ribbons (Uthaiah and Hudspeth, 2010b; Kantardzhieva et al., 2012), suggesting CaMKII may regulate RBC synaptic vesicle dynamics. Of note, protein kinase C α (PKCα), a well-known marker for these neurons (Zhang and Yeh, 1991; Kolb, Zhang and Dekorver, 1993; Casini et al., 1996; Kosaka et al., 1998; Caminos et al., 2001; Huh, Choi and Jeon, 2015), was expressed at a higher level than CaMKII in zebrafish RBC (Supplementary Fig. 1). In-situ hybridization chain reaction (HCR) with zebrafish CaMKII isoform-specific primers in retinas from the *Tg(vsx1:mem-Cerulean)* line, which highlights RBCs, largely corroborated the scRNA-seq analysis, where CaMKIIγ was most prevalent, followed by the δ2 isoform, then the β1 and β, and with minimal levels of the α (Figure 1).

**Figure 1.**
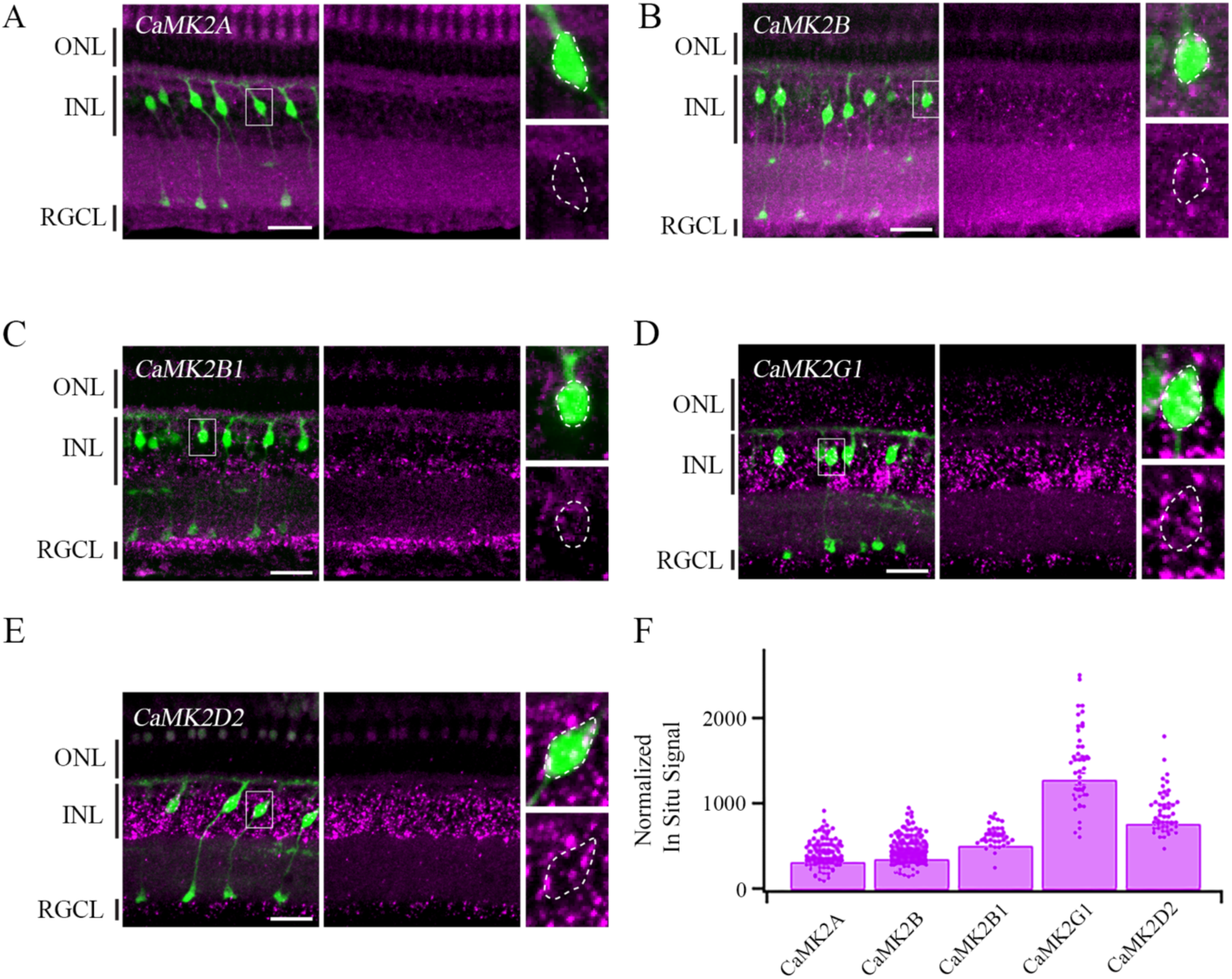
Expression patterns of CaMKII Isoforms. A-E. In situ Hybridization Chain Reaction (HCR) targeting CaMKII isoforms (magenta) in retinal cross sections of adult (9-12 months old) *Tg(vsx1:mem-Cerulean)* transgenic fish, which labels RBCs (green). **Left panel** shows a merged image of both HCR signal (magenta) and mem-Cerulean expression (green). The white rectangles indicate the cells shown in the high-magnification images on the right panels. **Middle panel** shows the HCR signal alone. **Right panels** show high-magnification views of the boxed regions shown in the left panel. Dashed lines outline the soma of mem-Cerulean-positive RBC1s. ONL, outer nuclear layer; INL, inner nuclear layer; RGCL, retinal ganglion cell layer. *CaMK2A,* CaMKIIα*; CaMK2B,* CaMKIIβ*; CAMK2B1,* CaMKIIβ1*; CaMK2G1,* CaMKIIγ1; *CaMK2D2,* CaMKIIδ2. **F.** Quantification of in situ HCR data. Bar graph showing the normalized in situ signal (see Materials and Methods) for the different CaMKII isoforms. Individual superimposed points show individual readings from different RBC somas. CaMKIIγ1 was the most prevalent, followed by CaMKIIδ2, then CaMKIIβ1 and CaMKIIβ, with minimal levels of CaMKIIα (χ²(4) = 218.7, p = 2.2e-12, Kruskal-Wallis test followed by Dunn’s post hoc test; CaMKIIβ vs. CaMKIIα p = 0.086, CaMKIIβ vs. CaMKIIβ1 p = 0.0001, CaMKIIβ vs. CaMKIIγ1 p = 5.08e-27, CaMKIIβ vs. CaMKIIδ2 p = 1.68e-17, CaMKIIβ1 vs. CaMKIIα p = 6.28e-7, CaMKIIβ1 vs. CaMKIIγ1 p = 9.9e-6, CaMKIIβ1 vs. CaMKIIδ2 p = 0.0088, CaMKIIγ1 vs. CaMKIIα p = 1.81e-30, CaMKIIδ2 vs. CaMKIIα p = 9.81e-21, CaMKIIδ2 vs. CaMKIIγ1 p = 0.11; N: CaMKIIα n = 103 cells, 3 fish, CaMKIIβ n = 149 cells, 3 fish, CaMKIIβ1 n = 37 cells, 1 fish, CaMKIIγ1 n = 46 cells, 2 fish, CaMKIIδ2 n = 48 cells, 2 fish).

### Reagents to acutely manipulate the level of CaMKII activity in rod bipolar cells

Previous studies investigating CaMKII’s roles in RBC synaptic transmission have reported conflicting findings, with one study suggesting that CaMKII facilitates synaptic vesicle release (Campbell et al., 2020), while another indicates little to no effect on exocytosis (Liang et al., 2021) in the mouse retina. This discrepancy may partly stem from differences in the specificity of the inhibitors, KN-62 and AIP, used in these studies to manipulate CaMKII activity. Indeed, KN-62 has been reported to have off-target effects (Sihra and Pearson, 1995; Marley and Thomson, 1996; Gargett and Wiley, 1997; Humphreys et al., 1998; Ledoux, Chartier and Leblanc, 1999; Brooks and Tavalin, 2011). AIP, and other inhibitors derived from the CaMKII autoinhibitory region, while tending to be more potent and selective than KN-62, have also been linked with off-target effects (Backs et al., 2009; Wu et al., 2009).

CaMKIIN is a natural CaMKII inhibitor protein expressed in the brain, which inhibits Ca^2+^/CaM-stimulated and autophosphorylation-generated Ca^2+^/CaM-independent (i.e. autonomous) CaMKII activity (Chang, Mukherji and Soderling, 1998, 2001), which contrast with that of KN-62, which only inhibits Ca^2+^/CaM-stimulated CaMKII activity. The inhibitory action from this protein was initially determined to arise from a 27-residue long segment of the protein, and an inhibitory peptide derived from this region was designated as CaMKIINtide (Chang, Mukherji and Soderling, 1998). Refinement of the minimal inhibitory determinants of CaMKIINtide and subsequent systematic substitutions within its sequence yielded an optimized 19-amino acid long peptide referred to as CN19o (Coultrap and Bayer, 2011). CaMKIINtide and CN19o have been used in multiple studies probing CaMKII contribution to various physiological processes, and thus far appear to be the most potent and selective inhibitors of CaMKII activity. CaMKIINtide, and presumably CN19o, bind within a continuous groove that connects the substrate binding site and the autophosphorylation/regulatory domain (Özden et al., 2022). CaMKII in general, and this binding region in particular, is highly conserved across isoforms (Tobimatsu and Fujisawa, 1989). As such, it would appear that CN19o may be useful to target all CaMKII isoforms. Indeed, CaMKIINtide and a subsequent shorter derivative, known as CN21, similarly inhibit CaMKIIα and CaMKIIβ (Chang, Mukherji and Soderling, 1998, 2001; Vest et al., 2007). Kinase activity assays further support this idea as Cy5-conjugated CN19o, at concentrations > 100 nM, virtually eliminated kinase activity from all human CaMKII isoforms tested, while a control Cy5-conjugated scrambled version of the peptide (CN19scr) showed only modest inhibition (Figs. 2A-C). In these assays, the Cy5-conjugated CN19o inhibited CaMKIIα with a similar IC50 to that originally reported (Coultrap and Bayer, 2011), suggesting that the fluorescent tag did not interfere with its ability to inhibit CaMKIIα (Fig. 2A). CaMKIIγ (Fig. 2B) and CaMKIIδ (Fig. 2C) exhibited weaker IC50’s than CaMKIIα. Additionally, CaMKIIδ displayed an apparent increased Hill co-efficient (Fig. 2C). Whether these differences in CN19o-mediated inhibition of CaMKII isoforms arise from their distinct association domains, which mediate holoenzyme assembly, is not clear. As all zebrafish CaMKII isoforms show a high (> 90%) conservation with their human counterparts (Rothschild, Lister and Tombes, 2007), and the zebrafish CaMKIIN is nearly identical to the rat CaMKIIN in the region from which CaMKIINtide and CN19o is derived (Hsu and Tseng, 2010), it appears that CN19o should be useful for examining the contribution of CaMKII to synaptic vesicle release in zebrafish. To support this idea, the interaction of CN19o with the kinase domain of zebrafish CaMKIIγ was modeled based on the previously determined structure of CaMKIINtide with human CaMKIIα. Examination of these structures suggests that CN19o adopts a conformation within the zebrafish CaMKII substrate binding pocket analogous to that of CaMKIINtide with human CaMKIIα (Fig. 2D). Collectively, from these data, we expect that when delivered to zebrafish RBC presynaptic terminals, at 1 μM, CN19o should ablate the majority of CaMKII activity.

**Figure 2.**
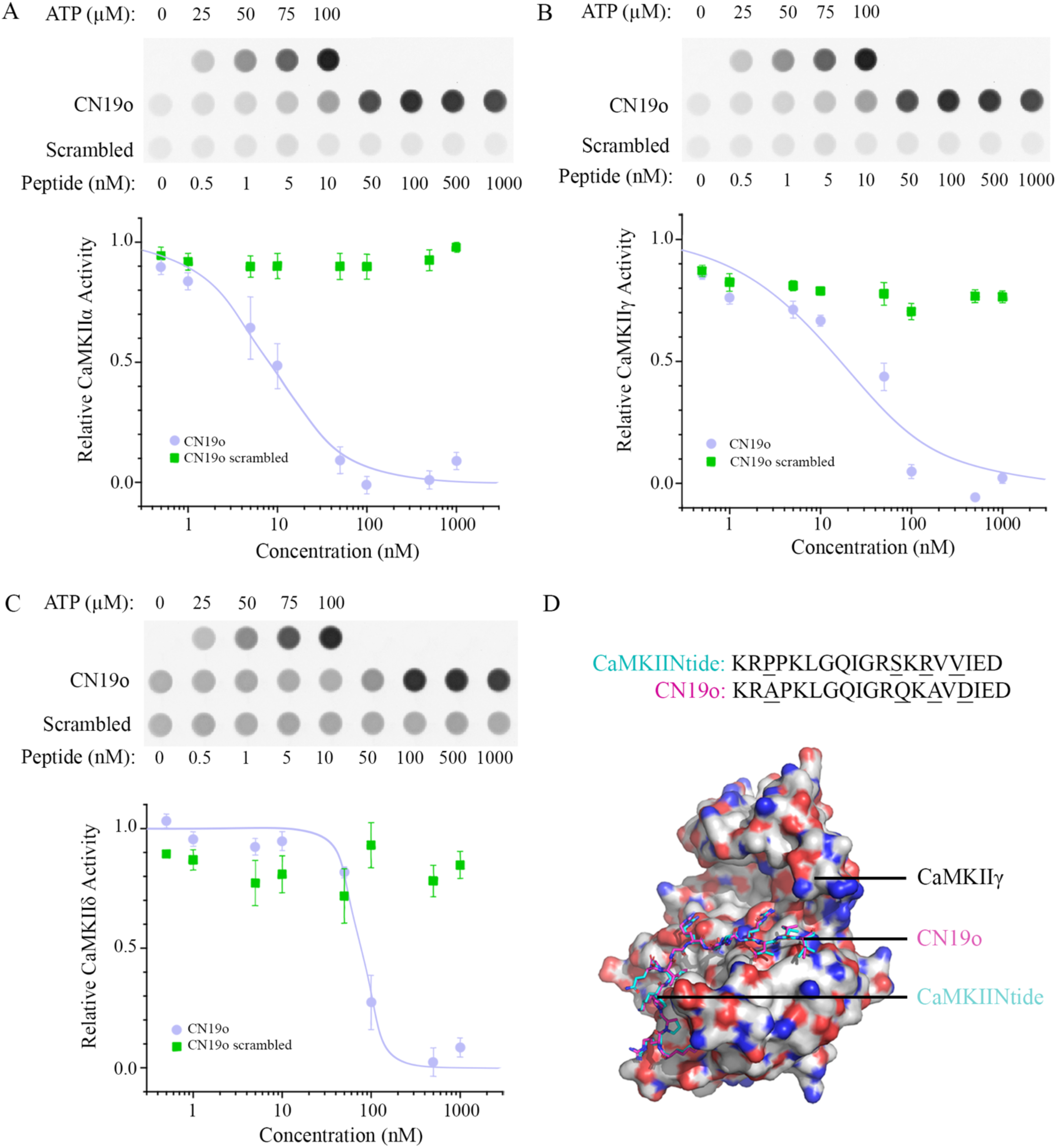
CN19o inhibition of CaMKII isoforms. Kinase assays were performed for **A.** CaMKIIα, **B.** CaMKIIγ, and **C.** CaMKIIδ. For each panel (**A-C**) ***Upper*** panels show ATP standard curve along with kinase assays performed in presence of CN19o or CN19scr peptide. ***Lower*** panel show the corresponding summary graph of CaMKII isoform activity (mean ± SEM) at different concentrations of CN19o (lilac) and CN19o scrambled (green) (n = 3). Kinase activity was determined relative to the signal acquired in the absence of peptides. A logistic fit to the CN19o data is shown. **D.** Interaction of CN19o with CaMKIIγ. **Upper.** The CN19o peptide alignment with CaMKIINtide with mutated residues underlined. **Lower.** Pymol (Molecular Graphics System, Version 1.2r3pre, Schrödinger, LLC) three-dimensional figure showing the superposition of CN19o (pink) and CaMKIItide (cyan) peptides and placement when interacting with the zebrafish CaMKIIγ isoform (shown in white, blue, and red).

An additional means to control CaMKII activity is to limit the availability of CaM, which is required for its Ca^2+^-dependent activation. Although KN-62 and KN-93 have long been thought to inhibit CaMKII activation by competing with Ca^2+^/CaM (Tokumitsu et al., 1990; Sumi et al., 1991), multiple off-target effects have been associated with these compounds (Brown and Bayer, 2024). Indeed, additional effects of these compounds likely arise from their direct interaction with CaM in either a Ca^2+^-independent (Brooks and Tavalin, 2011) or Ca^2+^-dependent (Wong et al., 2019) manner. The CaMBD of the SK2 Ca^2+^-activated K^+^ channel binds CaM constitutively with high affinity, irrespective of the presence of Ca^2+^ (Xia et al., 1998; Schumacher et al., 2001). As such, in some experiments, we took advantage of this feature and deployed a recombinant CaMBD (Supp. Fig. 2) as a method to buffer cellular CaM upon its infusion into RBC terminals and as an additional means to limit CaMKII activity.

By itself, acute application of CaMKII inhibitors can provide a gauge of the requirement of CaMKII in a physiological process such as transmitter release. However, this type of manipulation does not address the dynamic range available to the kinase. Thus, as a counterpart to our use of the CN19o and CaMBD peptides that limit CaMKII activity, our studies make use of a Ca^2+^-independent constitutively active (CA) his-tagged form of CaMKII (Hanson et al., 1989; Tavalin, 2008; Tavalin and Colbran, 2017) which, when acutely delivered to RBC terminals, should elevate CaMKII activity and drive phosphorylation of potential targets controlling synaptic vesicle release. Indeed, this truncated CA CaMKII exhibits Ca^2+^-independent kinase activity, which is largely ablated when heat inactivated (HI) (Supp. Fig. 3). As such, this latter HI preparation can serve as an appropriate control for the CA CaMKII.

### CaMKII inhibition reduces depolarization-evoked release

CaMKII has been long thought to facilitate vesicle release at various synapses (Llinás et al., 1985; Nichols et al., 1990; Jin and Hawkins, 2003; Lu and Hawkins, 2006; Pang et al., 2010), yet, mixed findings have been reported in mouse RBC ribbon synapses (Liu, Heidelberger and Janz, 2014; Campbell et al., 2020; Liang et al., 2021). Using zebrafish as a model organism, we examined whether CaMKII controls depolarized evoked vesicle release in RBCs by use of fluorescently labelled CN19o and its scrambled control. Neither reagent appeared to affect the calcium current amplitude at-10 mV compared to cells recorded without these peptides, and all showed overlapping current-voltage relationships with no changes in the Ca^2+^ current amplitude and charge (data not shown), suggesting that CaMKII does not affect depolarization-induced Ca^2+^ entry (Fig. 3B). These findings parallel those reported in mouse RBC using AIP to inhibit CaMKII (Campbell et al., 2020). Despite the similar calcium current evoked in response to a 250 ms stimulation (see Materials and Methods) in each of these conditions, the ensuing increase in membrane capacitance, reflective of vesicle release, was selectively reduced by 69.50% in cells dialyzed with CN19o compared to control whereas the CN19o scrambled peptide was similar to control (Fig. 3C, D). Thus, acute interference with CaMKII activity decreases evoked release at zebrafish RBC ribbon synapses, consistent with interference with a facilitatory action of CaMKII on exocytosis (Llinás et al., 1985; Nichols et al., 1990; Jin and Hawkins, 2003; Lu and Hawkins, 2006; Liu, Heidelberger and Janz, 2014). Further, we compared the release efficiency, defined as the change in capacitance over the integrated charge transfer (ι1Cm/Q_Ca_) between different groups and found that there is no difference between the control and the CN19o scrambled peptide, but CN19o substantially reduced the release efficiency, further supporting the idea that CaMKII tunes Ca^2+-^dependent vesicle release (Fig. 3E). Since the capacitance difference between CN19o scrambled and control conditions was similar, we compared CN19o to control in the remaining experiments.

**Figure 3.**
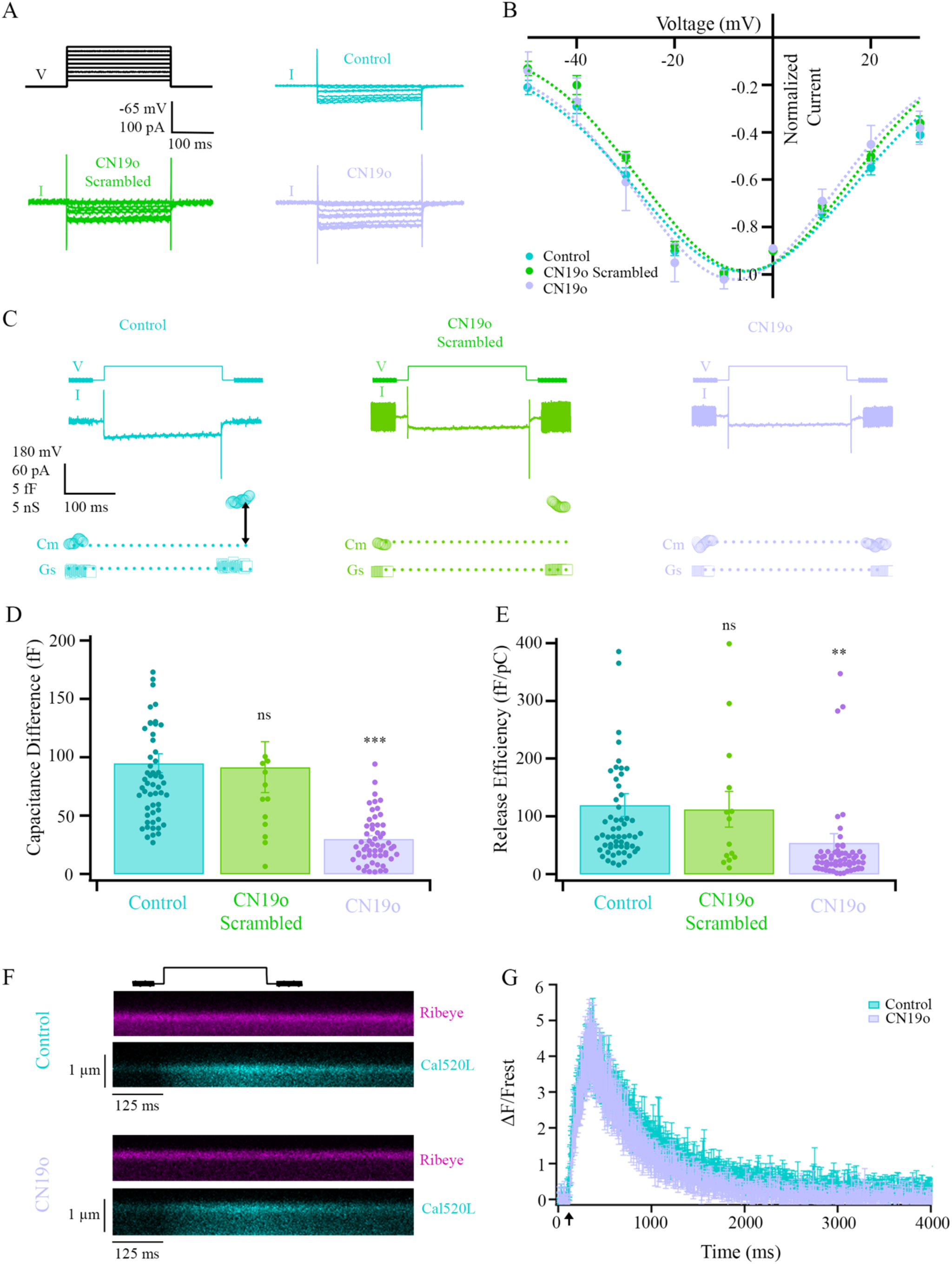
CN19o reduced vesicle fusion evoked by depolarization. **A.** An example of voltage steps (V) and current response (I) for control (cyan), CN19o scrambled (green), and CN19o (lilac). **B.** The plot illustrates the voltage-dependence of the average calcium current obtained from RBCs 5 minutes after delivering control pipette solution (cyan), CN19o scrambled (green), and CN19o (lilac) to the terminal via patch-pipette. CN19o does not alter the I-V relationship compared to control and CN19o scrambled. Vertical lines show ± SEM. The mean calcium current amplitude at-10 mV was similar across conditions (Control =-65.03 ± 8.63 pA; CN19o =-74.46 ± 12.15 pA; CN19o scrambled =-75.84 ± 13.06 pA), (χ²(2) = 0.81, p = 0.67, Kruskal-Wallis test followed by Dunn’s post hoc test; N: Control n = 6 fish, 9 cells; CN19o scrambled n = 5 fish, 7 cells; CN19o n = 4 fish, 5 cells). **C.** Sample of calcium current response to a 250 ms pulse in a cell dialyzed with control pipette solution (cyan) or with CN19o scrambled (green) and CN19o (lilac). Small sinusoidal stimuli used to monitor membrane capacitance (*C*_m_) and series conductance (*G*_s_) are present at the beginning and end of the voltage trace. The double arrow indicates the change in *C*_m_ evoked by the stimulus. **D.** Comparison of capacitance difference (Δ*C*_m_) from 14-62 synaptic terminals. Superimposed points show the individual readings for each condition. Results reveal that delivery of CN19o results in a significant decrease in capacitance response compared to control, whereas there were no significant differences between control and CN19o scrambled control (χ²(2) = 61, p = 7.3e-14, Kruskal-Wallis test followed by Dunn’s post hoc test; control vs. CN19o p = 9.37e-14, control vs. CN19o scrambled p = 0.46, CN19o vs. CN19o scrambled p = 0.00016; control n = 43 fish, 58 cells; CN19o n = 35 fish, 62 cells; CN19o scrambled n = 12 fish, 14 cells). ns = not significant, ***, p < 0.001. **E.** Comparison of release efficiency obtained by dividing the capacitance difference by the cathodic charge of individual synaptic terminals. Superimposed points show the individual measurements for each condition. Results reveal that delivery of CN19o significantly decreases release efficiency compared to control, whereas there were no significant differences between control and CN19o scrambled control (χ²(2) = 42, p = 7.9e-10, Kruskal-Wallis test followed by Dunn’s post hoc test; control vs. CN19o p = 7.4e-10, control vs. CN19o scrambled p = 0.47, CN19o vs. CN19o scrambled p = 0.0029; control n = 43 fish, 58 cells; CN19o n = 35 fish, 62 cells; CN19o scrambled n = 12 fish, 14 cells). ns, not significant; **, p < 0.005. **F.** Line scans were performed at a ribbon location to obtain synaptic Ca^2+^ responses elicited by 250-ms depolarization. The x-t plots show RBP-TAMRA fluorescence for ribbon location (top, magenta) and local Ca^2+^ signal fluorescence measured by ribbon-bound calcium indicator Cal520LA-RBP in control (cyan) and 5 mins after dialyzing CN19o (lilac). The extracellular space is the darker region at the top of each plot. The black trace on top of the line scans shows the timing of depolarization. **G.** Plot quantifying the average Cal520LA-RBP fluorescence (± SEM) as a function of time at zebrafish RBC individual ribbon location following a 250 ms stimulus for control (cyan) and CN19o (lilac) (control n = 10 cells, 7 fish; CN19o n = 7 cells, 5 fish). Peak Δ*F*/*F*_rest_ for control was 5.28 and for CN19o was 5.72. No significant differences were observed between the peaks of the two conditions (t(15) =-0.29, p = 0.78, two-sample t-test, control n = 10 cells, 7 fish; CN19o n = 7 cells, 5 fish). The black arrow shows the timing of 250 ms depolarization.

Though our data shows the calcium current was unaltered with CN19o, it remains unclear whether the reduction in the synaptic vesicle fusion is associated with altered ribbon-associated Ca^2+^ signals. To measure ribbon-associated Ca^2+^ signals, we conjointly delivered, via a whole-cell patch pipette, the Ca^2+^ indicator dye, Cal520L, conjugated to RBP (Cal520L-RBP) and a red-fluorescent dye conjugated to RBP (RBP-TAMRA), to RBC terminals (Vaithianathan and Matthews, 2014; Rameshkumar et al., 2024). Using this strategy, RBP-TAMRA effectively marks ribbons, while the Cal520L-RBP measures Ca^2+^ at the ribbon. Fluorescent intensity vs. time measurements reveal that the fluorescent intensity selectively rises in the Cal-520 channel upon membrane depolarization when cells were infused with our standard pipette solution or those in which CN19o was included (Fig. 3F). CN19o did not alter the normalized Ca^2+^ signal (Δ*F*/*F*_rest_) due to depolarization (Fig. 3G). Therefore, CaMKII inhibition by CN19o does not alter the overall Ca^2+^ signal or the Ca^2+^ signal in the vicinity of ribeye-associated synaptic vesicle pool. As such, the reduction in depolarization-evoked release, evaluated by capacitance differences, upon CaMKII inhibition is most likely due to processes downstream of calcium entry.

It is well-established that RBC ribbon synapses have kinetically distinct vesicle pools, including the ultrafast releasable pool (UFRP), the readily releasable pool (RRP), the recycling pool (RP), and the reserve pool (Res. P.) that are believed to be linked to distinct morphological correlates on the ribbon and within the cell (Fig. 4A, green, yellow, orange, and blue vesicles, respectively) (Mennerick and Matthews, 1996; von Gersdorff and Matthews, 1997; Neves and Lagnado, 1999; Zenisek, Steyer and Almers, 2000; Singer and Diamond, 2003; Joselevitch and Zenisek, 2020). Based on pool sizes and proximity to the membrane, UFRP vesicles are believed to be the population of vesicles docked at the base of the synaptic ribbon and contribute to the rapid first phase of neurotransmitter release. RRP vesicles that make up the second phase of release are believed to be the ones distal to the plasma membrane on the ribbon. The RP is believed to be the cytoplasmic pool that replenishes the ribbon pools and contributes to extrasynaptic release, and the Res. P. contains the cytoplasmic vesicles that do not participate in exocytosis. When brief pulses (0.5-10 ms) are applied to the RBC terminal, UFRP vesicles are released, leaving the RRP intact. As longer pulses are applied, more vesicles from the RRP begin to be used until the pool is depleted. To test the hypothesis that CaMKII controls the brief and sustained components of release, we measured the capacitance in response to various stimulus durations (10-1000 ms) in the absence and presence of CN19o (Fig. 4B, left panel). Capacitance measures were scaled to vesicle number by using a 26.4 aF/vesicle conversion factor, based on the ∼ 29 nm vesicle diameter in goldfish bipolar cells and a specific membrane capacitance is 10 fF/µm^2^ (Fettiplace, Andrews and Haydon, 1971; Breckenridge and Almers, 1987; von Gersdorff et al., 1996), to estimate the number of vesicles released (Fig. 4B, right panel). Based on an exponential fit, the initial rate constant of vesicle fusion was 470 fF/s under control conditions and was reduced to 140 fF/s in the presence of CN19o, corresponding to 18,077 and 5,385 vesicles/s, respectively. Despite the reduced number of vesicles release due to CaMKII inhibition, the time constants were slightly slower for control (∼302 ms) than in the presence of CN19o (∼215 ms), suggesting that in the face of continuous depolarization, CaMKII regulates the size of the releasable pool rather than inhibit the recruitment of new vesicles to replenish vesicles undergoing release. Of note, the number of vesicles fused following both brief (10 ms) and sustained pulses (above 100-ms) was larger in the control group compared to the RBC with CN19o, suggesting that CaMKII regulates the available pool size for both fast-transient and slow-sustained synaptic vesicle release. As vesicle fusion is compromised by CN19o even with the shortest duration stimuli, these data suggest that CaMKII, at minimum, likely acts at an early step required for vesicle release.

**Figure 4.**
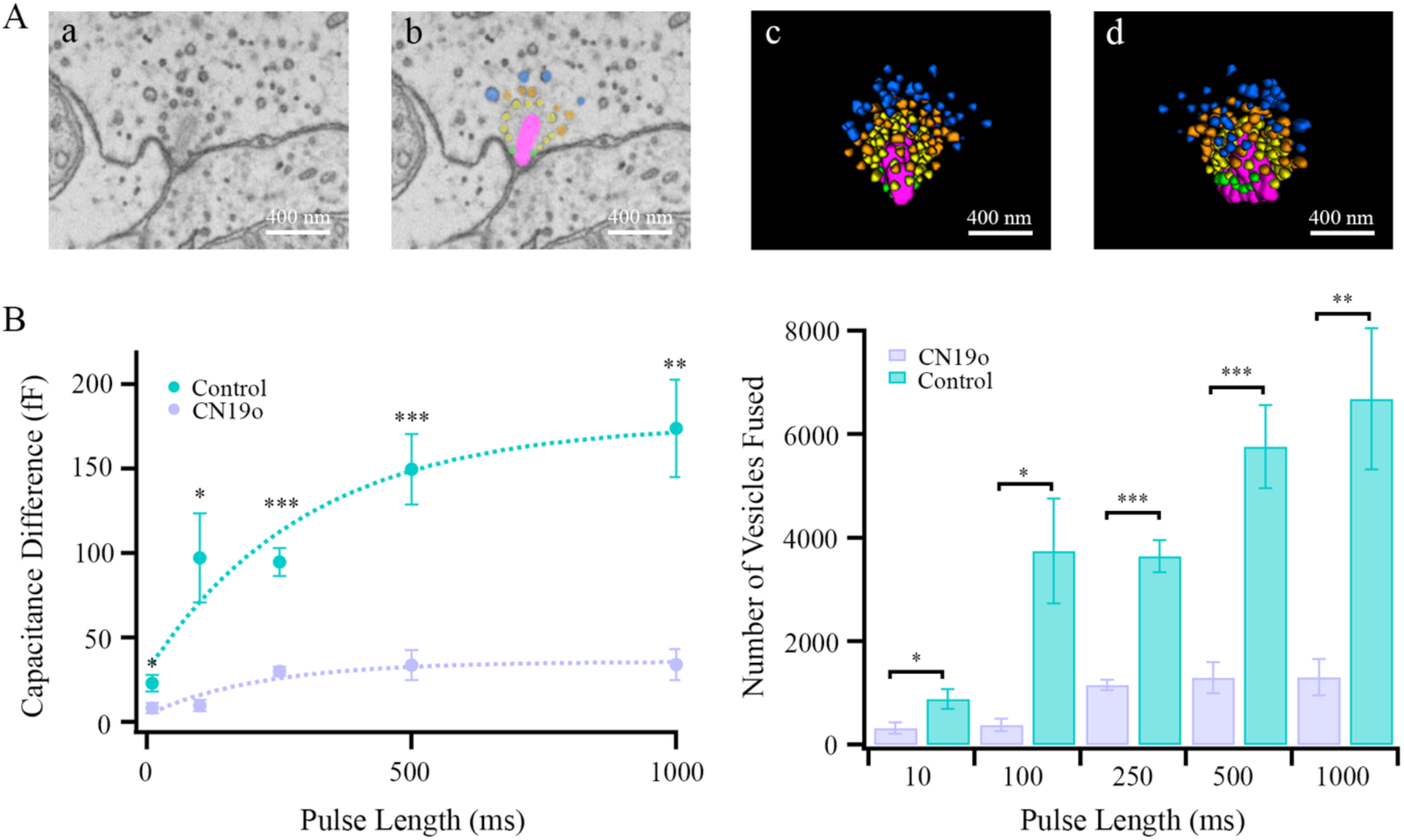
CN19o inhibits kinetically distinct synaptic vesicle pool release. A. **(a)** A single serial block-face scanning electron microscopy section shows a zebrafish RBC ribbon synapse. **(b)** Synaptic ribbons are highlighted in pink, and synaptic vesicle pools are color-coded based on kinetically distinct synaptic vesicle pools (green, vesicle pools at the base of the ribbon that likely comprise the ultrafast releasable pool; yellow, distal ribbon-associated pool; orange, recycle pool; blue, reserve pool). **(c-d)** 3D-rendered ribbon structure at two different angles for visualization of its synaptic vesicles. **B. Left Panel.** Plot summarizing the average increase in *C*_m_ (± SEM) evoked by voltage-clamp pulses of varying duration for control (cyan) and CN19o (lilac). Significant differences were found between control and CN19o in all pulse lengths measured (**10 ms:** t(18) = 2.68, p = 0.015, two-sample t-test, control n = 5 fish, 8 cells, CN19o n = 5 fish, 12 cells; **100 ms:** t(5.15) = 3.29, p = 0.021, Welch’s two-sample t-test, control n = 6 fish, 6 cells, CN19o n = 6 fish, 6 cells; **250 ms:** t(69.13) = 7.50, p = 0.00000000016, Welch’s two-sample t-test; control n = 43 fish, 58 cells; CN19o n = 35 fish, 62 cells; **500 ms:** t(7.62) = 5.23, p = 0.00093, Welch’s two-sample t-test, control n = 5 fish, 7 cells, CN19o n = 5 fish, 6 cells; **1000 ms:** t(7.17) = 4.63, p = 0.0022, Welch’s two-sample t-test, control n = 6 fish, 7 cells, CN19o n = 6 fish, 6 cells). Please note that the 250-ms data is the same as presented in **Figure 2D**. **Right Panel.** Visualization of the number of vesicles released with each pulse for control (cyan) and CN19o (lilac) based on data from **B**. *, p < 0.05; **, p < 0.01; ***, p < 0.001.

### CaMKII Inhibition Impairs Synaptic Vesicle Replenishment Following Longer Intervals

RBCs are specialized in transmitting information across multiple time frames, but with different outputs. Indeed, sustained depolarizations exhibit a decline in release, evaluated by capacitance measures, which reflect the depletion of the RRP (von Gersdorff and Matthews, 1997). However, this RRP can be replenished when synapses are given sufficient recovery time. The extent of synaptic depression and recovery can be evaluated by a two-pulse protocol, in which the interval between pulses is varied (von Gersdorff and Matthews, 1997). Given that CaM, CaMKII’s upstream activator protein, has been suggested to facilitate replenishment (Van Hook et al., 2014), we examined whether CaMKII contributes to either of these phenomena. A 250 ms depolarization suffices to deplete the RRP (Vaithianathan and Matthews, 2014). When, 250 ms depolarizations were evoked with a 10s interval between them, control cells and those in which CN19o was infused both exhibited a ∼ 60% reduction in the size of the second capacitance response (Fig. 5B). Thus, despite an overall reduction in the extent of release due to CaMKII inhibition (Fig. 3D and 4B), a similar fraction is replenished within a 10 s interval suggesting that CaMKII does not contribute to the early repopulation of the RRP. In contrast, when the interval between stimuli was 30 s, the second capacitance response fully recovered in control cells, but remained ∼ 50% depleted in cells with CN19o infused (Fig. 5B). The decrease in capacitance observed in the second pulse was not due to compromised Ca^2+^ entry as it was relatively stable irrespective of the interpulse interval for control cells and those infused with CN19o (Fig. 5C). Thus, CaMKII activity is required for full replenishment of the RRP, and this action is not secondary to changes in Ca^2+^ influx.

**Figure 5.**
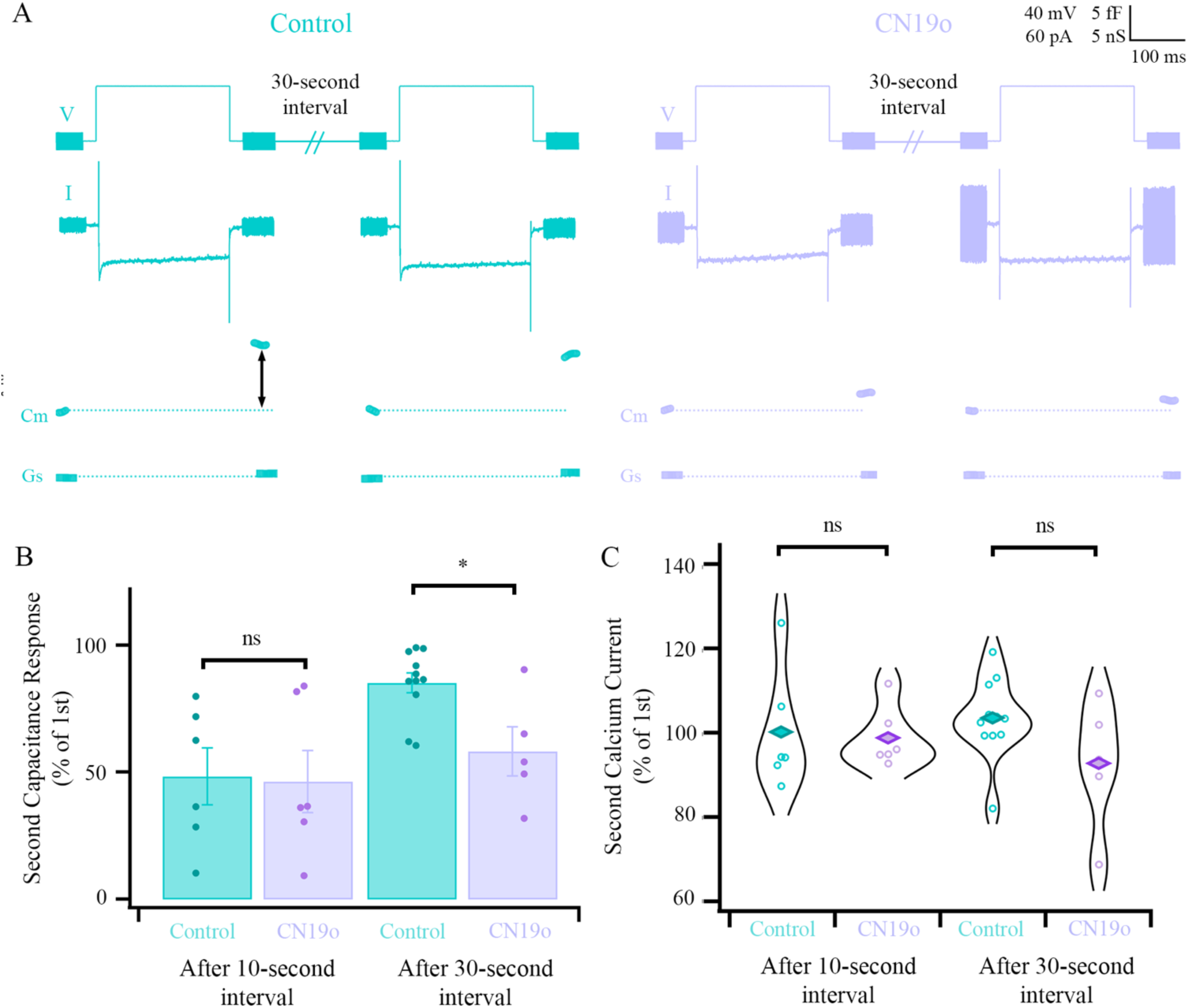
CN19o inhibits synaptic paired pulse recovery. **A.** Example of current responses (I), capacitance (*C*_m_), and series conductance (*G*_s_) in response to 250 ms paired-pulse stimulation with a 30s-interpulse interval (V), and for control (cyan) and CN19o (lilac). The double arrow indicates the change in *C*_m_ evoked by the stimulus. **B.** Summary of the experiments described in **A**. The second control Δ*C*_m_ and the Δ*C*_m_ in response to CN19o are expressed as a percentage of the first jump after 10-and 30-second intervals. Δ*C*_m_ in response to CN19o shows significant changes compared to control following a 30-second interval (t(14) = 3.12, p = 0.0075, two-sample t-test, control n = 10 fish, 11 cells; CN19o n = 4 fish, 5 cells) but not a shorter 10-second interval (t(10) = 0.12, p = 0.91, two-sample t-test, control n = 6 fish, 6 cells, CN19o n = 6 fish, 6 cells). **C.** The same experiments as in **B**, showing the corresponding calcium current amplitudes. The second calcium currents as a percentage of the first were not significantly different between control and CN19o following the 10-second (t(10) = 0.26, p = 0.80, two-sample t-test, control n = 6 fish, 6 cells, CN19o n = 6 fish, 6 cells) and 30-second intervals (t(14) = 1.79, p = 0.094, control n = 10 fish, 11 cells; CN19o n = 4 fish, 5 cells). ns, not significant, *, p < 0.05.

### Efficient Synaptic Vesicle Release Requires CaMKII Activation by CaM

Since Ca^2+^/CaM serves as the principal activating signal for CaMKII, we sought to determine if interfering with CaM availability would effectively mimic the effects of CaMKII inhibition by CN19o. Buffering cellular CaM levels by infusion of the recombinant SK2 channel-derived CaMBD (10 μM) into RBC terminals did not alter the peak Ca^2+^ current amplitude and did not produce any overt alterations in the I-V (Fig 6A). However, similar to CN19o, delivery of CaMBD resulted in a reduction in the depolarization-evoked capacitance difference (Fig. 6B), which was accompanied by a decrease in release efficiency (Fig. 6C). These data are consistent with a substantial portion of CaMKII’s contribution to release being at least CaM-and presumably Ca^2+^-dependent.

**Figure 6.**
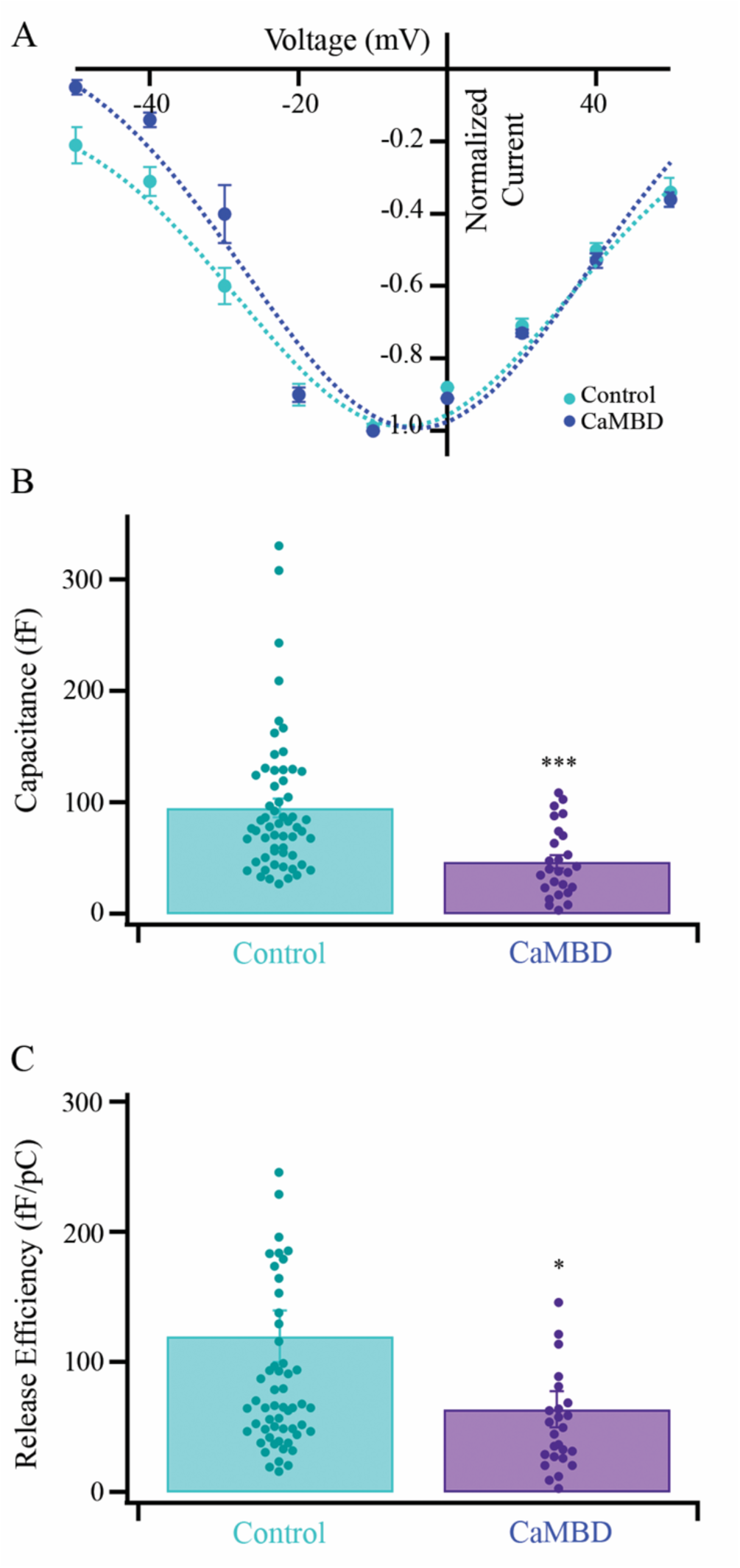
CaMBD inhibits vesicle fusion evoked by depolarization. **A.** The plot illustrates the voltage-dependence of the average calcium current normalized to the peak current in control (cyan) or 5 minutes after delivering of CaMBD (purple). The mean Ca^2+^ current amplitude at-10 mV was similar between conditions (Control =-65.03 ± 8.63 pA, CaMBD =-97.05 ± 10.40 pA), with no significant differences between them (t(11) = 2.00, p = 0.071, two-samples t-test, control n = 6 fish, 9 cells, CaMBD n = 4 cells, 3 fish). Please note that the control data is the same as shown in **Figure 2B**. **B.** Bar graph summarizing the Δ*C*_m_ evoked by 250 ms pulses in control (cyan) and CaMBD (purple). Superimposed points show individual readings for each condition. Δ*C*_m_ was significantly reduced with CaMBD delivery (t(80.9) = 4.71, p = 0.000010, Welch’s two-sample t-test, control n = 43 fish, 58 cells, CaMBD n = 11 fish, 26 cells). Please note that the control data is the same as shown in **Figures 2D** and **3B**. ***, p < 0.001. **C.** Comparison of release efficiency obtained by dividing the capacitance difference by the cathodic charge of individual synaptic terminals. Superimposed points show the individual measurements for each condition. Results reveal that delivery of CaMBD significantly decreases release efficiency compared to control (t(81.7) = 2.33, p = 0.022, Welch’s two-sample t-test, control n = 43 fish, 58 cells, CaMBD n = 11 fish, 26 cells). ns, not significant; *, p < 0.05.

### Constitutively Active CaMKII on Synaptic Vesicle Dynamics in Rod Bipolar Cell Ribbon Synapses

Given our observation that CaMKII inhibition decreases vesicle fusion and replenishment in RBC, we hypothesized that increasing active CaMKII levels in the RBC terminal would result in the opposite observations. For these experiments, we delivered a constitutively active (CA) Ca^2+/^CaM-independent CaMKII (Supp. Fig. 3) into terminals. The mean Ca^2+^ current amplitude at-10 mV at the peak of the I-V was similar to control conditions, independent of whether CA or the heat-inactivated (HI) control was infused into the terminals (Fig. 7A). In conjunction with our previous experiments using either CN19o or the CaMBD to suppress CaMKII activity, these data suggest that Ca^2+^ influx at zebrafish RBC terminals is relatively insensitive to the acute bidirectional manipulation of CaMKII activity.

**Figure 7.**
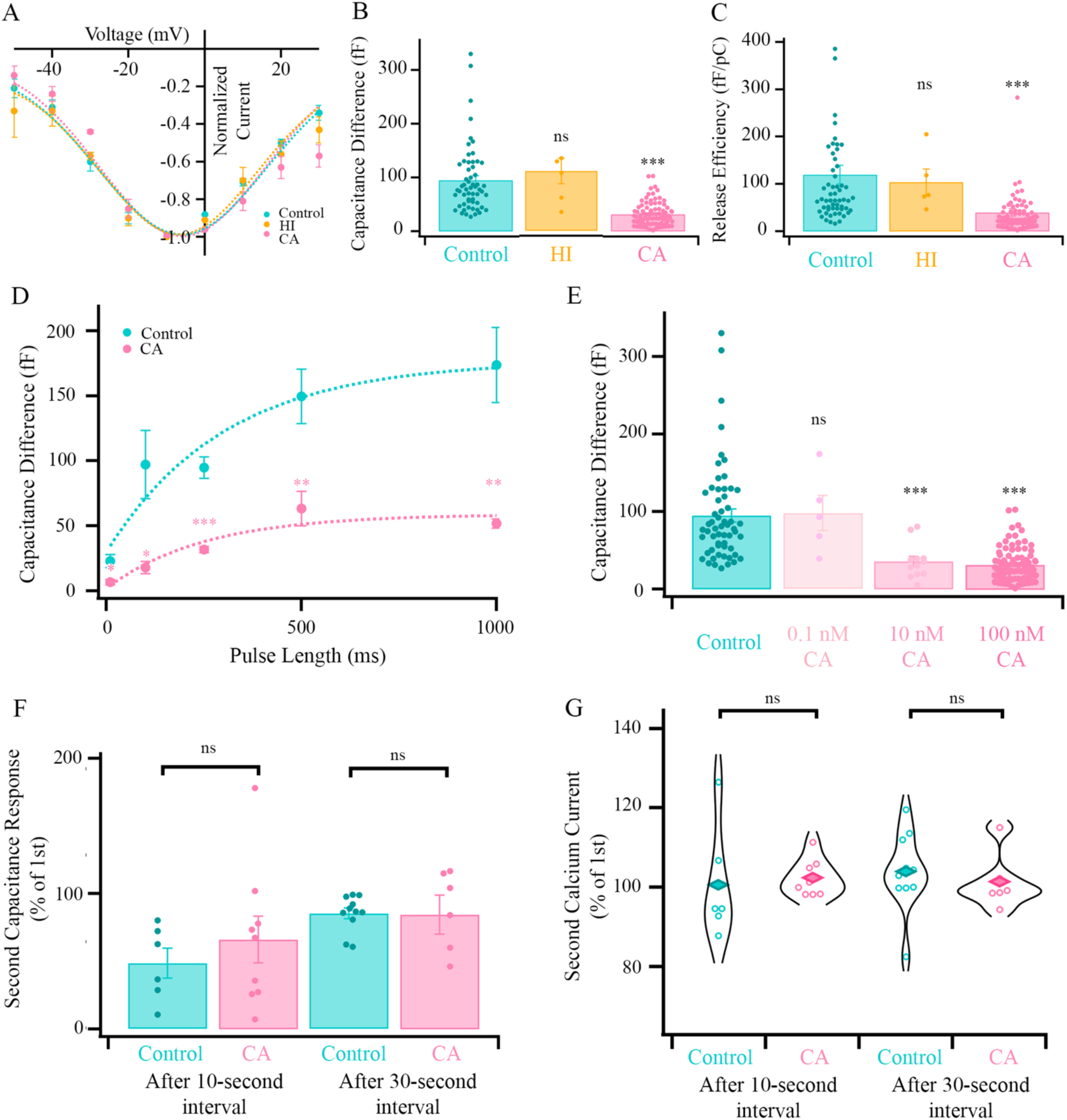
Delivery of constitutively active CaMKII to the RBC terminal shows distinct effects in release and replenishment. **A.** Summary of the normalized current-voltage relationship following delivery of 100 nM CA (pink), HI (orange), or control (cyan) to the RBC terminal. The mean Ca^2+^ current amplitude at-10 mV was similar across conditions (Control =-65.03 ± 8.63 pA; HI =-95.40 ± 18.82 pA; CA =-82.65 ± 15.42 pA), with no significant differences between them (F(2,13) = 1.5, p = 0.26, one-way ANOVA followed by Tukey’s HSD post hoc test; N: control n = 6 fish, 9 cells; HI n = 3 fish, 3 cells; CA n = 3 fish, 4 cells). Please note that the control data is the same as shown in **Figures 2B and 5B. B.** Comparison of capacitance difference (Δ*C*_m_) from 6-88 synaptic terminals. Superimposed circles show the individual points for each condition. Results reveal that delivery of CA results in a significant decrease in capacitance response compared to control, whereas there were no significant differences between control and HI control (χ²(2) = 70, p = 6.5e-16, Kruskal-Wallis test followed by Dunn’s post hoc test; control vs. CA p = 1.65e-15, control vs. HI p = 0.80, CA vs. HI p = 0.0024; control n = 43 fish, 58 cells, CA n = 45 fish, 88 cells, HI n = 5 fish, 5 cells). Please note that the 250 ms control data is the same as shown in **Figures 2D, 3B, and 5C**. ns = not significant, ***, p < 0.001. **C.** Comparison of release efficiency obtained by dividing the capacitance difference by the cathodic charge of individual synaptic terminals. Superimposed points show the individual measurements for each condition. Results reveal that delivery of CA significantly decreases release efficiency compared to control, whereas there were no significant differences between control and HI control (χ²(2) = 55, p = 9.2e-13, Kruskal-Wallis test followed by Dunn’s post hoc test; control vs. CA p = 3.3e-12, control vs. HI p = 0.57, CA vs. HI p = 0.0027; control n = 43 fish, 58 cells, CA n = 45 fish, 88 cells, HI n = 5 fish, 5 cells). ns, not significant; ***, p < 0.001. **D.** Graph summarizes the average Δ*C*_m_ in response to varying duration of stimuli for control (cyan) and CA (pink). Our data shows significant differences between control and CA in all pulse durations (**10ms:** t(9.20) = 3.01, p = 0.014, Welch’s two-sample t-test, control n = 5 fish, 8 cells, CA n = 4 fish, 6 cells; **100ms:** t(5.27) = 2.97, p = 0.029, Welch’s two-sample t-test, control n = 6 fish, 6 cells, CA n = 7 fish, 7 cells; **250ms:** t(66.35) = 7.37, p = 0.00000000034, Welch’s two-sample t-test, control n = 43 fish, 58 cells, CA n = 45 fish, 88 cells; **500ms:** t (11) = 3.37, p = 0.0063, two-sample t-test, control n = 5 fish, 7 cells; CA n = 6 fish, 6 cells; **1000ms:** t(6.17) = 4.21, p = 0.0053, Welch’s two-sample t-test, control n = 6 fish, 7 cells, CA n = 5 fish, 6 cells). Please note that the 250 ms control data is the same as shown in **Figures 2D, 3B, 5C, and 6B**. **E.** Bar graph summarizing the Δ*C*_m_ in response to varying concentrations of CA CaMKII. Control is shown in cyan, and the different concentrations of CA are shown in different shades of pink. Superimposed points indicate individual readings. Capacitance difference increased as the concentration of CA CaMKII delivered decreased, with significant differences between control and 100 nM CA and 10 nM CA, but no differences between control and 0.1 nM CA (χ²(3) = 73, p = 8.2e-16, Kruskal-Wallis test followed by Dunn’s post hoc test, control vs. 0.1 nM CA p = 0.80, control vs. 10 nM CA p = 0.00062, control vs. 100 nM CA p = 1.6e-15, 0.1 nM CA vs. 10 nM CA p = 0.037, 0.1 nM vs. 100 nM CA p = 0.004, 10 nM CA vs 100 nM CA p = 1.00; N: control n = n = 43 fish, 58 cells, 0.1 nM CA n = 4 fish, 5 cells, 10 nM CA n = 6 fish, 12 cells, 100 nM CA n = 45 fish, 88 cells). Please note that the 250 ms control data is the same as shown in **Figures 2D, 3B, 5C, 6B, and 6C**. **F.** The second control Δ*C*_m_ and the Δ*C*_m_ in response to CA CaMKII are expressed as a percentage of the first jump after 10-and 30-second intervals. Superimposed points show individual readings. Δ*C*_m_ was not significantly different between control and CA following the 10-(t(13) = 0.77, p = 0.455, two-sample t-test, control n = 6 fish, 6 cells, CA n = 8 fish, 9 cells) and 30-second intervals (t(6.12) =-0.20, p = 0.85, Welch’s two-sample t-test, control n = 10 fish, 11 cells; CA n = 6 fish, 6 cells). **G.** The same experiments as in E, showing the corresponding calcium current amplitudes. The second calcium currents as a percentage of the first were not significantly different between control and CA following the 10-(t(5.64) =-0.30, p = 0.77, Welch’s two-sample t-test, control n = 6 fish, 6 cells, CA n = 8 fish, 9 cells) and 30-second intervals (t(15) = 0.56, p = 0.58, two-sample t-test, control n = 10 fish, 11 cells; CA n = 6 fish, 6 cells). CA CaMKII, Constitutively Active CaMKII; HI, Heat-Inactivated CA CaMKII; ns, not significant; *, p < 0.05; **, p < 0.01; ***, p < 0.001.

Infusion of the HI CaMKII did not substantially alter the capacitance change induced by a 250 ms depolarization, whereas, contrary to our hypothesis, CA CaMKII infusion into RBC terminals decreased the capacitance change by ∼70% (Fig. 7B). Similarly, we also found there were no differences in the release efficiency between the RBC infused with control vs. HI, but a CA CaMKII reduced release efficiency compared to control (Fig. 7C) Since the capacitance difference between control and HI conditions was similar, we compared CA to control in the remaining experiments. Regardless of the stimulus duration, CA CaMKII decreased the capacitance change when compared to control (Fig. 7D). Exponential fits to plots of the capacitance change vs. stimulus duration were similar in the absence (∼302 ms) and presence of CA CaMKII (∼252 ms). In conjunction, with our data for CaMKII inhibition by CN19o (Fig. 4) these data suggest that the overall release kinetics were comparable, irrespective of CaMKII activity, but that pool size decreased with overactive or underactive CaMKII. We estimated the corresponding initial vesicle release rate to be 217.8 fF/s (or 8377 vesicles/s) in the presence of CA CaMKII. Given the deviation from our expectations, we infused lower concentrations of the CA CaMKII to ascertain whether there was a critical level at which elevation of CaMKII activity impairs synaptic release. Indeed, infusion of CA CaMKII at concentrations above 0.1 nM led to impairments of release (Fig. 7E). Given that either inhibition (Fig. 3) or elevation of CaMKII activity (Fig. 7) hinders release in the absence of any systematic changes in Ca^2+^ influx, these data collectively suggest that endogenous CaMKII likely exists at an optimal level to maintain exocytosis at ribbon synapses.

As CaMKII inhibition not only impaired vesicle release (Figs. 3 and 4) but also vesicle replenishment (Fig. 5), and elevation of CaMKII activity also impaired release (Figs. 7B and D), we examined whether elevated CaMKII activity would similarly impair vesicle replenishment using the two-pulse protocol. Here, infusion of CA CaMKII,did not alter the degree of capacitance reduction evident after a either a 10 s or 30 s inter-pulse interval (Fig. 7F). This dramatically contrasts with the impaired replenishment at 30 s observed with CaMKII inhibition with CN19o (compare with Fig. 5B). Importantly, this effect occurred in the absence of any alteration in the Ca^2+^ influx elicited with either a 10 s or a 30 s inter-pulse interval (Fig. 7G). Therefore, elevating CaMKII activity in RBC terminals through the delivery of CA CaMKII permitted full replenishment of the releasable pool despite the apparent reduced capacity (Fig. 7B). Collectively, these data suggest that while CaMKII is endogenously present at optimal levels to ensure release of the RRP, ongoing CaMKII activity is essential for ensuring the replenishment of this vesicle pool size regardless of its magnitude.

## Discussion

While all CaMKII isoforms are present and exhibit distinct distribution across retinal cell types (Sun et al., 2024), our scRNA-seq and HCR analysis of zebrafish RBCs reveals that CaMKIIγ and CaMKII8 are the predominant isoforms expressed at the transcript level in zebrafish. Whether this is also reflected at the protein level in zebrafish RBC terminals is unclear. However, CaMKII8 appears to be highly prevalent at these terminals in mice (Tetenborg et al., 2017). Thus, our data suggest that there may be some common utilization of distinct CaMKII isoforms to control synaptic transmission at RBC terminals across these species.

Given the potential presence of multiple CaMKII isoforms at zebrafish RBC terminals, we ensured that the CN19o peptide could inhibit all CaMKII isoforms. Indeed, CN19o virtually eliminated the activity of all isoforms at the concentration we eventually used for these studies. Although we did not directly evaluate CN19o against zebrafish CaMKII isoforms, our structural modeling suggested that it would interact with the most prevalent zebrafish CaMKII isoform in a manner highly analogous to that obtained with the human CaMKIIα and CaMKIINtide, from which CN19o is derived. Additionally, we took steps to ensure that a control scrambled version exhibited limited inhibitory activity against all CaMKII isoforms, thus bolstering confidence that effects due to CN19o could be attributed to its action as a CaMKII inhibitor. Using CN19o, our findings most closely align with those performed with AIP to inhibit CaMKII (Campbell et al., 2020). Indeed, CN19o (Figs. 3 and 4) and AIP (Campbell et al., 2020) inhibited release without notable effect on the triggering Ca^2+^ influx. It is well established that L-type calcium channels are the predominant source for triggering vesicle release from RBCs (Heidelberger and Matthews, 1992; Logiudice, Henry and Matthews, 2006). Our scRNA-seq data analysis of zebrafish retinal bipolar cells shows the presence of CaV1.3 and CaV1.4 calcium channels in RBCs, similar to the findings from mouse scRNA-seq (Shekhar et al., 2016). The lack of effect of CaMKII inhibition on Ca^2+^ influx is noteworthy as CaV1.3 calcium channels can be enhanced by CaMKII, but only in the presence of densin-180 (Jenkins et al., 2010). Our scRNA-seq data analysis of zebrafish retinal bipolar cells shows that *LRRC7,* a homolog of densin-180, is present at very low levels in RBCs (Supplementary Fig. 4), possibly indicative its preferential presence at postsynaptic sites (Apperson et al. 1996). Whether any CaMKII modulates CaV1.4 activity is unclear. As such, zebrafish RBC Ca^2+^ influx at synaptic terminals may be relatively resistant to acute manipulations in CaMKII activity. Consistent with this idea, neither the CaMBD nor the CA CaMKII had any appreciable effect on depolarization-induced Ca^2+^ current (Figs. 6A and 7A, respectively). Collectively, this suggests that the participation of CaMKII in the regulation of vesicle release from zebrafish RBCs is distal to Ca^2+^ entry, or the Ca^2+^ signal at the ribbon (Figs. 3F and G), and thus lies either in control of the fusion process and/or the mobilization and/or accessibility of the synaptic vesicles pool to the fusion machinery.

Our findings contrast with a recent study in which KN-62 was used to inhibit CaMKII, and W-7 and calmidazolium were used to antagonize CaM (Liang et al., 2021). As noted above, both methodological, species differences, and known off-target effects of these compounds, including actions on Ca^2+^ channels, could contribute to the discrepancy. An additional distinction between KN-62 and CN19o is that KN-62 would only be able to suppress Ca^2+^/CaM-stimulated CaMKII activity (Tokumitsu et al., 1990), while CaMKIINtide/CN19o and AIP inhibit Ca^2+/^CaM-dependent and Ca^2+^/CaM-independent CaMKII activity (Ishida and Fujisawa, 1995; Chang, Mukherji and Soderling, 1998). Thus, it is possible that a substantial fraction of the CaMKII activity in RBC terminals is acting in a Ca^2+^/CaM-independent manner. Indeed, autophosphorylated CaMKII, reflective of constitutively active Ca^2+^/CaM-independent CaMKII, is readily detected at hair cell ribbon synapses (Uthaiah and Hudspeth, 2010b). In line with this idea, we found that the depolarization-induced capacitance change was less sensitive to the CaMBD peptide relative to CN19o (Fig. 6B compare with Fig. 3D). The presence of an endogenous constitutively active CaMKII pool at RBC terminals might be anticipated given that they experience relatively continuous Ca^2+^ influx under physiological conditions. Of note, even autonomous CaMKII can be further subject to Ca^2+/^CaM stimulation (Barcomb et al., 2014), thereby providing an additional mechanism by which CaMKII activity can be fine-tuned across its dynamic range.

Our data suggest that an optimal level of CaMKII activity is required to support vesicle pool size (Figs. 4B and 7D) as either impairing or enhancing activity led to a reduction in overall release with minimal change in release kinetics. While our finding that elevating CaMKII activity, by introducing CA CaMKII to the RBC terminal, reduced vesicle release, was not expected, CaMKIIα has been reported to limit NTR at hippocampal synapses (Chapman et al., 1995; Hinds et al., 2003).

At present it is not clear how NTR is bidirectionally coupled to CaMKII activity levels. One possibility is that multiple CaMKII substrates exist, with some favoring and others limiting NTR. Not mutually exclusive with this scenario is the possibility that some CaMKII substrates may undergo successive cycles of phosphorylation and dephosphorylation during NTR, such that driving either phosphorylation or dephosphorylation hinders progression of NTR. Indeed, a similar scenario involving phosphorylation cycling has been proposed to control specific pools of postsynaptic receptors (Summers, Bogard and Tavalin, 2019). In either case, it will be important to identify countervailing phosphatases that either limit basal phosphorylation or restore the basal phosphorylation of those CaMKII substrates coordinating NTR. Although the focus of this study is on the action of CaMKII on NTR from zebrafish RBCs, it is important to note that other Ca^2+^-and/or Ca^2+^/CaM-dependent kinases such as PKC and MLCK have also been implicated in the control of transmitter release from ribbon synapses (Midorikawa et al., 2007; Ruether et al., 2010; Liang et al., 2021). To ascertain their respective roles, it will be critical to decipher whether these kinases share substrates sites with each other or whether they phosphorylate distinct sites on common and/or different substrates involved in NTR.

The critical CaMKII substrates that control NTR and their correspondence to distinct aspects of the neurotransmitter vesicle cycle remain unclear. While CaMKII, itself, is a likely substrate (Giese et al., 1998; Uthaiah and Hudspeth, 2010b), syntaxin-3B, has also been identified CaMKII substrate, whose phosphorylation state appears correlated with NTR at these synapses (Curtis et al., 2010; Liu, Heidelberger and Janz, 2014; Campbell et al., 2020; Hays et al., 2020), numerous others likely exist. Indeed, a large array of proteins exist at retinal ribbon synapses (Uthaiah and Hudspeth, 2010b; Thoreson and Zenisek, 2024). The relative degree to which each of these proteins are essential as opposed to modulatory remains to be fully elucidated. Indeed, distinct synaptotagmin isoforms harbor different roles for distinct phases of release in RBCs (Mesnard et al., 2022b). In this regard, it is noteworthy that the effects of acute CaMKII inhibition on NTR, shown here, closely phenocopy those due to genetic removal of ribeye as both manipulations lead to a reduction in releasable pool size and recovery from paired-pulse depression, without altering the kinetics of release (Mesnard et al., 2022a). Together, these findings suggests that ribeye function may be under the direct control of CaMKII activity and/or that ribeye controls CaMKII localization and/or activity. Determining the relevant CaMKII substrates contributing to distinct steps of vesicle docking, mobilization, fusion, and recycling represents a major challenge. Delineation of those substrates that contribute to the CaMKII regulation of NTR may enable the development of therapeutic strategies to mitigate maladaptive CaMKII signaling, suggested to participate in various retinopathies (Sun et al., 2024).

---

**Supplementary Figure 1.**
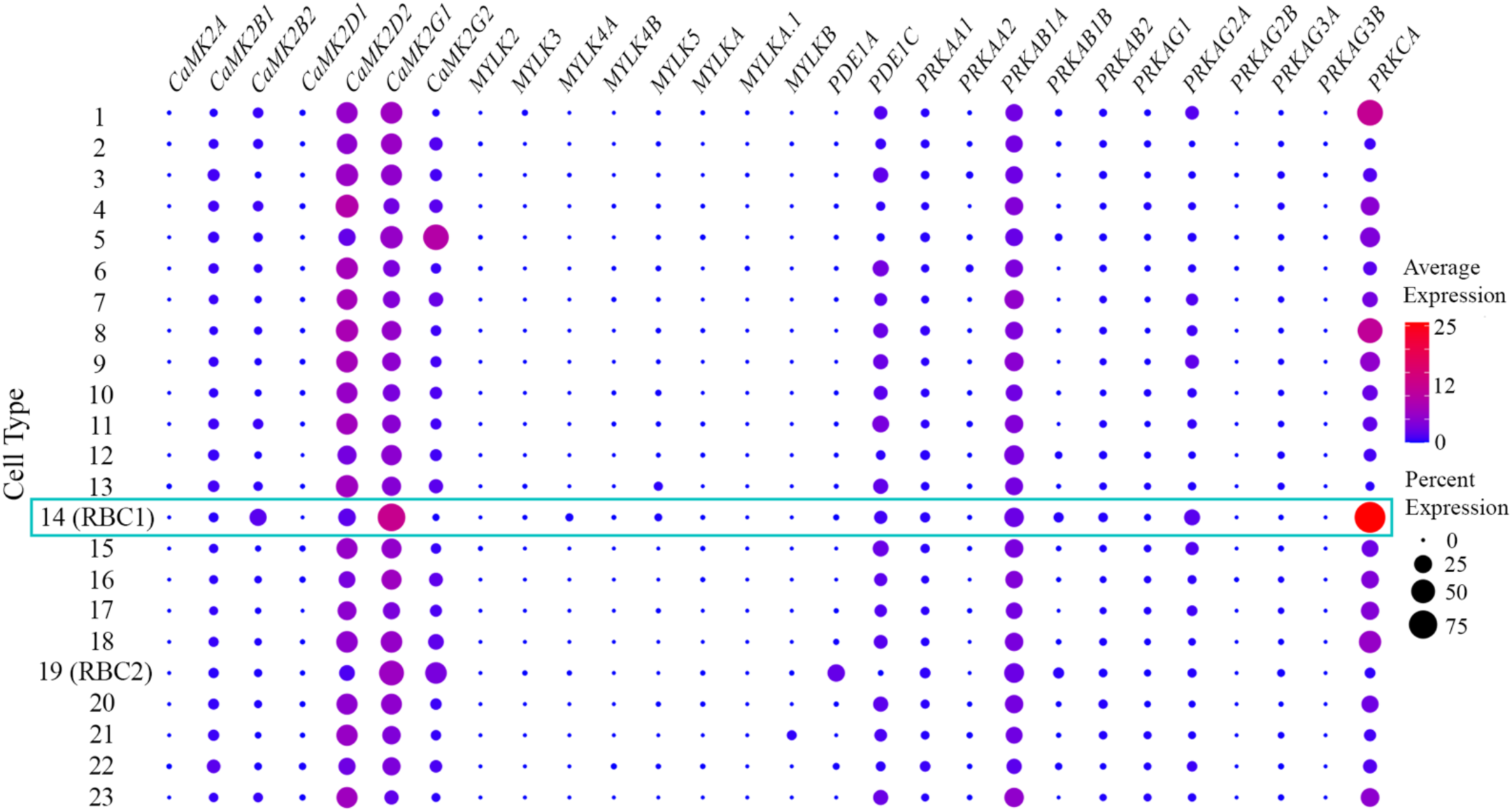
Single-cell RNA sequence analysis reveals differential gene expression patterns of CaM-activated kinases in zebrafish bipolar cells. The protein that the gene encodes is given at the top, and the bipolar cell types are given in the left column. Our RBC of interest, the RBC1, is indicated by the number 14 and marked with a cyan rectangle. Each dot in the plot indicates the average expression and percentage of expressing cells for a specific gene in a particular cell type. The size of the dots corresponds to the percentage of cells in the cluster in which the gene was detected, with larger dots indicating a higher percentage of expression. The color of the dots represents the average expression level of the gene, using a gradient that ranges from blue (low expression) to red (high expression). The x-axis represents different CaMKII genes, while the y-axis represents cell types, with 21 cone bipolar cells and 2 RBCs (numbered 14, RBC1; and 19, RBC2). The predominant isoform detected in both RBC types is γ1. In RBC1, the order of isoform prevalence is as follows: γ1 > δ2 > β2 > β1 > γ2 > α1 > δ1, with negligible levels of γ2, α1, and δ1. In RBC2, the prevalence order is γ1 > γ2 > δ2 > β1 > β2 > δ1 > α1, with minimal levels of β2, δ1, and α1. Myosin light-chain kinase (MYLK), phosphodiesterase 1 (PDE1), and AMP-activated protein kinase (AMPK, shown as PRKA above) levels in zebrafish RBCs are smaller than CaMKII’s. Protein kinase C α (PKCα, shown as PRKC above) expression in zebrafish RBCs is the highest. Genes list: *CaMK2A*, CaMKIIα; *CaMK2B1*, CaMKIIβ1; *CAMK2B2*, CaMKIIβ2; *CaMK2D1*, CaMKIIδ1; *CaMK2D2*, CaMKIIδ2; *CaMK2G1*, CaMKIIγ1; *CaMK2G2*, CaMKIIγ2; *MYLK2*, myosin light chain kinase 2; *MYLK3*, myosin light chain kinase 3; *MYLK4A*, myosin light chain kinase family, member 4a; *MYLK4B*, myosin light chain kinase family, member 4b; *MYLK5*, myosin light chain 5; *MYLKA*, myosin light chain a; *MYLKA.1*, myosin light chain a.1; *MYLKB*, myosin light chain b; *PDE1A*, phosphodiesterase 1A; *PDE1C*, phosphodiesterase 1C; *PRKAA1*, AMPKα1; *PRKAA2*, AMPKα2; *PRKAB1A*, AMPKβ1 non-catalytic subunit a; *PRKAB1B*, AMPKβ1 non-catalytic subunit b; *PRKAB2*, AMPKβ2; *PRKAG1*, AMPKγ1; *PRKAG2A*, AMPKγ2 non-catalytic subunit a; *PRKAG2B*, AMPKγ2 non-catalytic subunit b; *PRKAG3A*, AMPKγ3 non-catalytic subunit a; *PRKAG3B*, AMPKγ3 non-catalytic subunit b; *PRKCA*, PKCα.

**Supplementary Figure 2.**
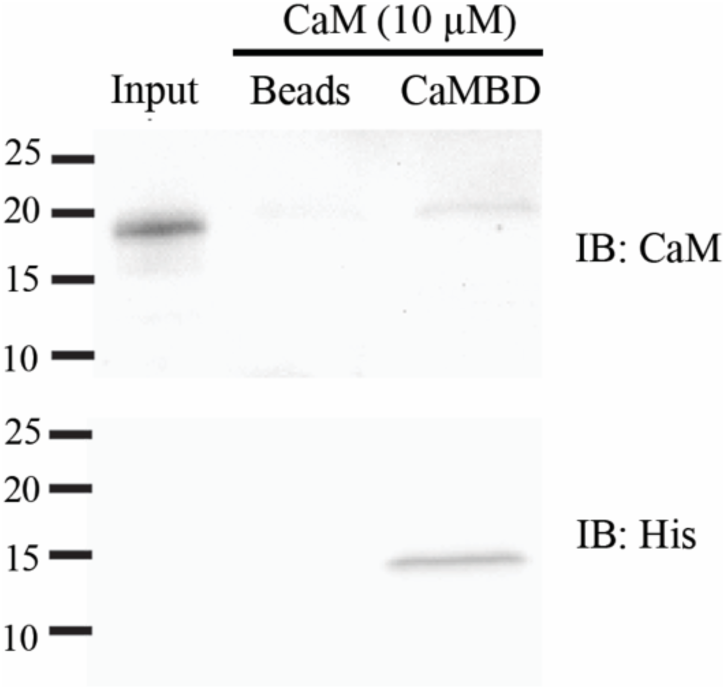
CaMBD Assay. CaM binding by CaMBD peptide. Ni-NTA beads or Ni-NTA beads coupled to His-CaMBD peptide (∼ 5 mg) were incubated with CaM (10 µM). **Upper.** Representative western blot showing the amount of CaM recovered by the peptide or beads alone control. A CaM standard is shown representing ∼0.1% of input. **Lower.** Blots were stripped and reprobed with anti-His HRP antibodies to reveal the presence of the CaMBD peptide.

**Supplementary Figure 3.**
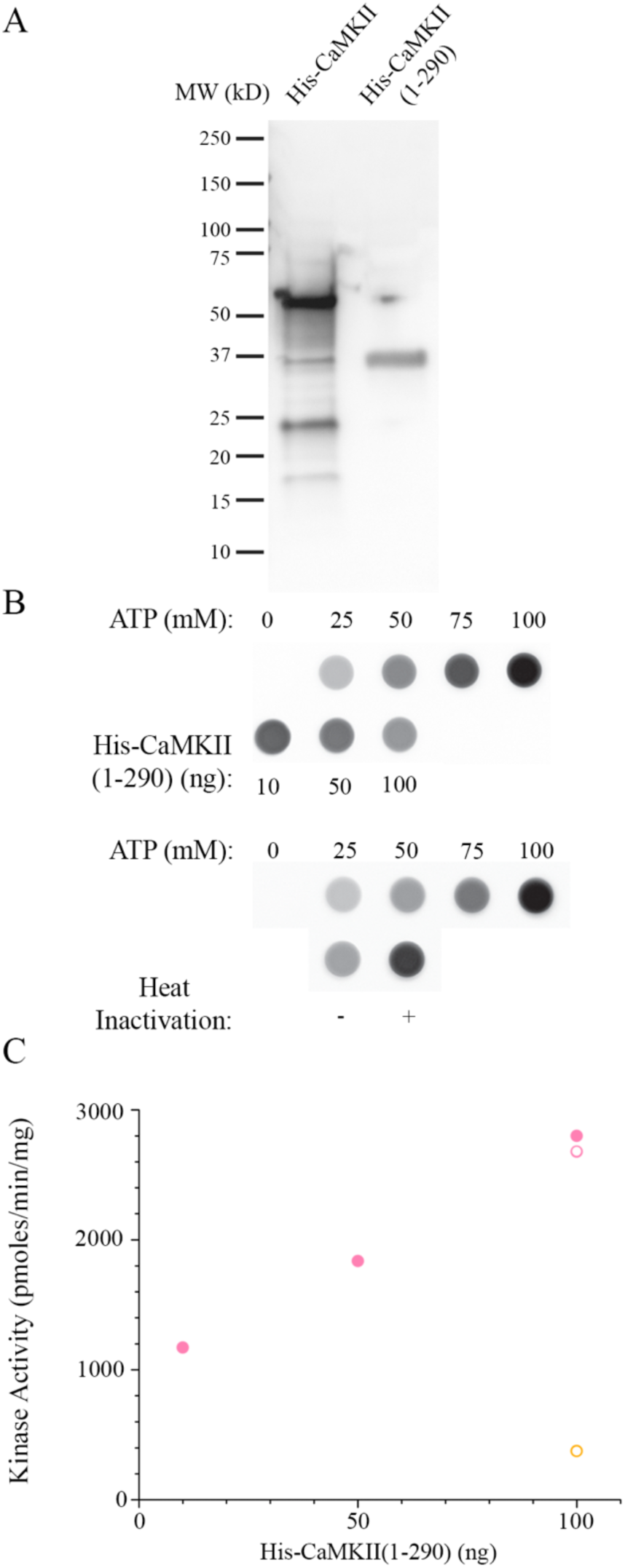
Purification and verification of constitutive His-CaMKIIα(1-290) activity. **A.** Representative western blot of commercially available full-length His-CaMKIIα vs His-CaMKIIα(1-290). Gels were loaded with 2 μg protein and probed with an anti-His HRP antibody. His-CaMKIIα(1-290) migrates at ∼37 kDa consistent with its predicted MW. **B.** Kinase assays performed in the absence of Ca^2+^/CaM. For each assay an ATP standard curve (0-100 μM) was run alongside it calibrate the amount of ATP consumed by the CaMKII reaction. **Upper.** Kinase activity with varying amounts of His-CaMKIIα(1-290). **Lower.** Comparison of kinase activity for His-CaMKIIα(1-290) and heat-inactivated His-CaMKIIα(1-290) using 100 ng of each in the assay. **C.** Quantified specific activity for assays shown in **B**. Filled circles represent the activity for the assay in the top panel of **B**. Open circles represent the activity for the bottom panel with the heat-inactivated His-CaMKIIα(1-290) in orange and the active His-CaMKIIα(1-290) in pink. In each assay, kinase activity was determined relative to the signal acquired at 100 mM ATP in the absence of kinase.

**Supplementary Figure 4.**
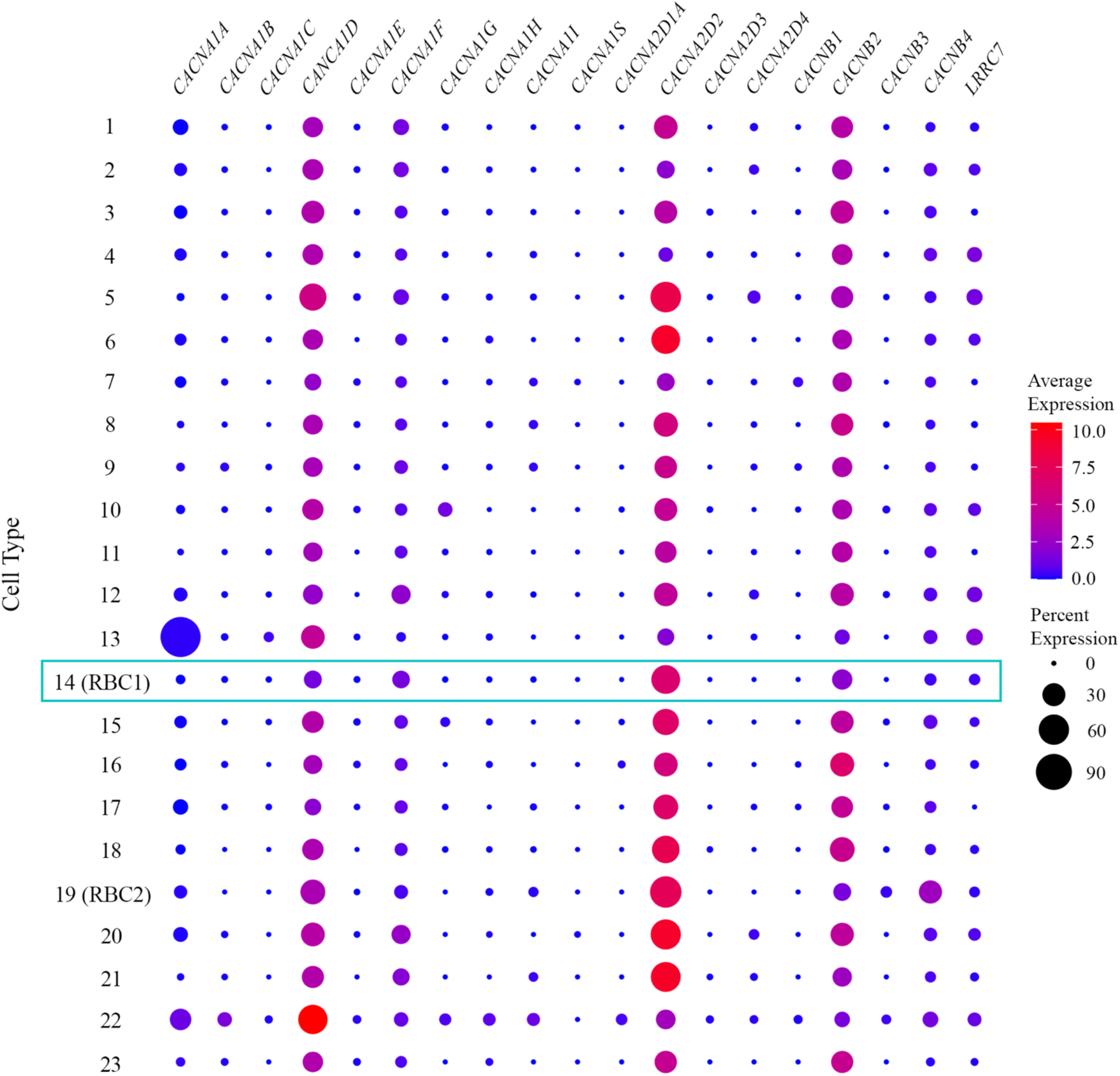
Single-cell RNA sequence analysis of calcium channels and dynosin-180 in zebrafish bipolar cells. The protein that the gene encodes is given at the top, and the bipolar cell types are given in the left column. Our RBC of interest, the RBC1, is indicated by the number 14 and marked with a cyan rectangle. Each dot in the plot indicates the average expression and percentage expression of a specific gene in a particular cell type. The size of the dots corresponds to the percentage expression of the gene, with larger dots indicating a higher percentage expression. The color of the dots represents the average expression level of the gene, using a gradient that ranges from blue (low expression) to red (high expression). The x-axis represents different genes, while the y-axis represents cell types, with 21 cone bipolar cells and 2 RBCs (numbered 14, RBC1; and 19, RBC2). Genes list: *CACNA1A,* CaV2.1*; CACNA1B,* CaV2.2*; CACNA1C,* CaV1.2*; CACNA1D,* CaV1.3*; CACNA1E,* CaV2.3*; CACNA1F,* CaV1.4*; CACNA1G,* CaV3.1*; CACNA1H,* CaV3.2*; CACNA1I,* CaV3.3*; CACNA1S,* CaV1.1*; CACNA2D1A,* CaVα281*; CACNA2D2,* CaVα282*; CACNA2D3,* CaVα283*; CACNA2D4,* CaVα284*; CACNB1,* CaVβ1*; CACNB2,* CaVβ2*; CACNB3,* CaVβ3*; CACNB4,* CaVβ4; LRRC7, dyosin-180.

