## Supplementary material for "Distinct Roles of CaMKII in Synaptic Vesicle Dynamics at Zebrafish Retinal Rod Bipolar Ribbon Synapses": Table 1

| | | |
|---|---|---|
| **CamKIIalpha probe set** | **1** | 5'- cctggaagcgtgtgcgcctcttctg TAgAAgAgTCTTCCTTTACg -3' |
| | **2** | 5'- gAggAgggCAgCAAACgg AA tccgcgacgagatgtgaagattccc -3' |
| | **3** | 5'- cccctcaacctcaatggcgaggcca TAgAAgAgTCTTCCTTTACg -3' |
| | **4** | 5'- gAggAgggCAgCAAACgg AA atctgccaacttcacagctgcgcct -3' |
| | **5** | 5'- caggtctacagccttcccataggga TAgAAgAgTCTTCCTTTACg -3' |
| | **6** | 5'- gAggAgggCAgCAAACgg AA cttccggaggacttcaggggacagg -3' |
| | **7** | 5'- cctcaggagtgactgtgtcccactc TAgAAgAgTCTTCCTTTACg -3' |
| | **8** | 5'- gAggAgggCAgCAAACgg AA gtgaaggaaaatcataggcgccggc -3' |
| | **9** | 5'- ctcctcgccgatcagatggatgtga TAgAAgAgTCTTCCTTTACg -3' |
| | **10** | 5'- gAggAgggCAgCAAACgg AA gttgaggatggtagtgtgaaccggc -3' |
| | **11** | 5'- acgatgagaagctgggagcttctgc TAgAAgAgTCTTCCTTTACg -3' |
| | **12** | 5'- gAggAgggCAgCAAACgg AA cctcaaacgactgatcctgcgactg -3' |
| | **13** | 5'- caaagatgaggtagtggtggccttc TAgAAgAgTCTTCCTTTACg -3' |
| | **14** | 5'- gAggAgggCAgCAAACgg AA ctgagatgctgtcatgcagacgcac -3' |
| | **15** | 5'- tgtttgagcgcctccgcagcagtga TAgAAgAgTCTTCCTTTACg -3' |
| | **16** | 5'- gAggAgggCAgCAAACgg AA cgtttggacgggttgatggtcagca -3' |
| | **17** | 5'- gattcgtgtttcttcggactgggca TAgAAgAgTCTTCCTTTACg -3' |
| | **18** | 5'- gAggAgggCAgCAAACgg AA gcagggcatcccgttattgtctagg -3' |
| | **19** | 5'- tctgcagatcctagcctccctgtcc TAgAAgAgTCTTCCTTTACg -3' |
| | **20** | 5'- gAggAgggCAgCAAACgg AA cttctggtggtcccgagtagacagc -3' |

| | | |
|---|---|---|
| **CamKIIbeta probe set** | **1** | 5'- tcgtcggtgaagcgcgtgcaggttg TAgAAgAgTCTTCCTTTACg -3' |
| | **2** | 5'- gAggAgggCAgCAAACgg AA gttgccatgtctgtcctcttccagc -3' |
| | **3** | 5'- cgtattcctgacccgtgctgagttt TAgAAgAgTCTTCCTTTACg -3' |
| | **4** | 5'- gAggAgggCAgCAAACgg AA cacatcggcgcaccacggagaaagc -3' |
| | **5** | 5'- atggaggcgcacgatgttggagtgt TAgAAgAgTCTTCCTTTACg -3' |
| | **6** | 5'- gAggAgggCAgCAAACgg AA tagcagacggcagatccgcgcctca -3' |
| | **7** | 5'- aagcaccgcctccaggatctgctgg TAgAAgAgTCTTCCTTTACg -3' |
| | **8** | 5'- gAggAgggCAgCAAACgg AA gcaatgactggcgtccgcttcactg -3' |
| | **9** | 5'- tgtagaggatgactccacacgccca TAgAAgAgTCTTCCTTTACg -3' |
| | **10** | 5'- gAggAgggCAgCAAACgg AA tgtccaccggcttcccatacgcttc -3' |
| | **11** | 5'- cacggtggagcgttggcagacccac TAgAAgAgTCTTCCTTTACg -3' |
| | **12** | 5'- gAggAgggCAgCAAACgg AA gtgtttgagagcctcctgcgctgtg -3' |
| | **13** | 5'- gcgcgacaccagcatggtggtcagt TAgAAgAgTCTTCCTTTACg -3' |
| | **14** | 5'- gAggAgggCAgCAAACgg AA ggctcccttgagtttccgtctggcg -3' |
| | **15** | 5'- gccatccacaaactgtgtcaggcgg TAgAAgAgTCTTCCTTTACg -3' |
| | **16** | 5'- gAggAgggCAgCAAACgg AA gtaggcgatgcaggcgccatcttca -3' |
| | **17** | 5'- gactatcggcttctaggctggagtt TAgAAgAgTCTTCCTTTACg -3' |
| | **18** | 5'- gAggAgggCAgCAAACgg AA cagagtgctggcgtcccattggtcc -3' |
| | **19** | 5'- caccaggttccccagagcctcgggt TAgAAgAgTCTTCCTTTACg -3' |
| | **20** | 5'- gAggAgggCAgCAAACgg AA gaaagacgtcagccctgggtcacac -3' |

| | | |
|---|---|---|
| | **1** | 5'- ccttgattcggcgagctggagcgtc TAgAAgAgTCTTCCTTTACg -3' |
| | **2** | 5'- gAggAgggCAgCAAACgg AA ggatattcgcctcgacgttgagctc -3' |
| | **3** | 5'- cacacagcacatgtgcacgacaagg TAgAAgAgTCTTCCTTTACg -3' |
| | **4** | 5'- gAggAgggCAgCAAACgg AA agtctctgcacgggtggaaagacac -3' |

| | | |
|---|---|---|
| CamKIIbeta1 probe set | **5** | 5'- ggtttaatcgagtctgggcgctttc TAgAAgAgTCTTCCTTTACg -3' |
| | **6** | 5'- gAggAgggCAgCAAACgg AA aacaccactggctcagctgaacacc -3' |
| | **7** | 5'- gacgccagtcactgcatccagcaga TAgAAgAgTCTTCCTTTACg -3' |
| | **8** | 5'- gAggAgggCAgCAAACgg AA ctggaggcggtgcttcactgccatc -3' |
| | **9** | 5'- gtctccctgcacctcaatggccagt TAgAAgAgTCTTCCTTTACg -3' |
| | **10** | 5'- gAggAgggCAgCAAACgg AA aaagtcggccagcttcacagcagcg -3' |
| | **11** | 5'- ctcctgtctgtgcatcatggaggcc TAgAAgAgTCTTCCTTTACg -3' |
| | **12** | 5'- gAggAgggCAgCAAACgg AA agtggagcgttggcagacccatggg -3' |
| | **13** | 5'- ctcatcctccactgtggcattactg TAgAAgAgTCTTCCTTTACg -3' |
| | **14** | 5'- gAggAgggCAgCAAACgg AA gtccgacgactcctttacgtctgct -3' |
| | **15** | 5'- tccaccaggtttcccaaggcctcag TAgAAgAgTCTTCCTTTACg -3' |
| | **16** | 5'- gAggAgggCAgCAAACgg AA tcaaagcaggtgagtccagggtcgc -3' |
| | **17** | 5'- tccaatcaggtgcacgtgagggttg TAgAAgAgTCTTCCTTTACg -3' |
| | **18** | 5'- gAggAgggCAgCAAACgg AA gatggtggtgtggatgggcttactg -3' |
| | **19** | 5'- ggtcggccctggccatccacatact TAgAAgAgTCTTCCTTTACg -3' |
| | **20** | 5'- gAggAgggCAgCAAACgg AA gtgaggcggatgtaggcaatgcagg -3' |
| CamKIIdelta2 probe set | **1** | 5'- tgatgatgcggggaagcgcagtcag TAgAAgAgTCTTCCTTTACg -3' |
| | **2** | 5'- gAggAgggCAgCAAACgg AA tcctcactcgtccctcctgtagacc -3' |
| | **3** | 5'- gccgcatactcttggccggatgata TAgAAgAgTCTTCCTTTACg -3' |
| | **4** | 5'- gAggAgggCAgCAAACgg AA ttcacgcatcgcctcacgacagaga -3' |
| | **5** | 5'- acgacaaatcctggcctctctctcc TAgAAgAgTCTTCCTTTACg -3' |
| | **6** | 5'- gAggAgggCAgCAAACgg AA cttctggtgatccctggcggacagc -3' |
| | **7** | 5'- ccccttcatcttactggccaacagc TAgAAgAgTCTTCCTTTACg -3' |
| | **8** | 5'- gAggAgggCAgCAAACgg AA attctcaggcttcaggtccctgtgg -3' |
| | **9** | 5'- cctggagtaccagcaaatccaaacc TAgAAgAgTCTTCCTTTACg -3' |
| | **10** | 5'- gAggAgggCAgCAAACgg AA gcctgctgatcaccctggacctcaa -3' |
| | **11** | 5'- gtttctgtccactgggttgtggatc TAgAAgAgTCTTCCTTTACg -3' |
| | **12** | 5'- gAggAgggCAgCAAACgg AA ggtagtttgtggctccagtgctggg -3' |
| | **13** | 5'- gcggatgtacgcgatacaggccgca TAgAAgAgTCTTCCTTTACg -3' |
| | **14** | 5'- gAggAgggCAgCAAACgg AA ctccccaatcaggtgcacatgaggg -3' |
| | **15** | 5'- atctcgacggtgccagacacgggtc TAgAAgAgTCTTCCTTTACg -3' |
| | **16** | 5'- gAggAgggCAgCAAACgg AA ctccgactgcatggtgcgtggcatc -3' |
| | **17** | 5'- ccagcagaatgtagaggatcacacc TAgAAgAgTCTTCCTTTACg -3' |
| | **18** | 5'- gAggAgggCAgCAAACgg AA aagcccacatgtccaccggtttgcc -3' |
| | **19** | 5'- gcactccacggtttcctgtctgtgc TAgAAgAgTCTTCCTTTACg -3' |
| | **20** | 5'- gAggAgggCAgCAAACgg AA cattgaggcaacagtggagcgttgg -3' |
| ıa1 probe set | **1** | 5'- ccctttccgagctcttcgtaaagct TAgAAgAgTCTTCCTTTACg -3' |
| | **2** | 5'- gAggAgggCAgCAAACgg AA tactcgtccgtaaacctggtcgagg -3' |
| | **3** | 5'- cccagcaaaaccaaaccaggcctgc TAgAAgAgTCTTCCTTTACg -3' |
| | **4** | 5'- gAggAgggCAgCAAACgg AA atctccctgcacctcgatggcaagc -3' |
| | **5** | 5'- ccagatgtccacaggtttgccgtag TAgAAgAgTCTTCCTTTACg -3' |
| | **6** | 5'- gAggAgggCAgCAAACgg AA atccttcctcaacacctctggagac -3' |
| | **7** | 5'- cgtgtcccactccggagaaggaaaa TAgAAgAgTCTTCCTTTACg -3' |
| | **8** | 5'- gAggAgggCAgCAAACgg AA ataagctccggccttgatctgctgg -3' |
| | **9** | 5'- gtgcatcatggacgccactgtggac TAgAAgAgTCTTCCTTTACg -3' |
| | **10** | 5'- gAggAgggCAgCAAACgg AA ttggcagacccatgggtgcttgagg -3' |

| | | |
|---|---|---|
| CamKIIgamr | 11 | 5'- ggcactgcactgactcatcacggca TAgAAgAgTCTTCCTTTACg -3' |
| | 12 | 5'- gAggAgggCAgCAAACgg AA gtcactgctcgctccctgcgcactt -3' |
| | 13 | 5'- cagcgtcctctccaatcagatgaac TAgAAgAgTCTTCCTTTACg -3' |
| | 14 | 5'- gAggAgggCAgCAAACgg AA gaggattgaggatagtggtgtgcac -3' |
| | 15 | 5'- gaatccaggcaagctgtcctgtgcg TAgAAgAgTCTTCCTTTACg -3' |
| | 16 | 5'- gAggAgggCAgCAAACgg AA tcactgtaatggggcagctggtgcc -3' |
| | 17 | 5'- ggaaaaggacaaaacagcaccgcca TAgAAgAgTCTTCCTTTACg -3' |
| | 18 | 5'- gAggAgggCAgCAAACgg AA tgggcgtcagatacatgcaccgtcc -3' |
| | 19 | 5'- cggggttgtgcacaactgtagtttg TAgAAgAgTCTTCCTTTACg -3' |
| | 20 | 5'- gAggAgggCAgCAAACgg AA gctccatcggtgtagaagaggccgt -3' |

| | | |
|---|---|---|
| Amplifier | B1-1 | 5'-CgTAAAggAAgACTCTTCCCgTTTgCTgCCCTCCTCgCATTCTTTCTTgA |
| | B1-2 | [ATTO647NN]-5'-gAggAgggCAgCAAACgggAAgAgTCTTCCTTTACgCTCTT |

ggAgggCAgCAAACgggAAgAg –3’-[ATTO647NN]
CCCgTTTgCTgCCCTCCTCAAgAAAgAATgC-3’
